# Mass spectrometric characterisation of the circulating peptidome following oral glucose ingestion in control and gastrectomised patients

**DOI:** 10.1101/2020.03.23.002899

**Authors:** Richard G. Kay, Rachel E. Foreman, Geoff P. Roberts, Richard Hardwick, Frank Reimann, Fiona M. Gribble

## Abstract

Meal ingestion triggers secretion of a variety of gut and endocrine peptides, several of which are routinely measured in research studies by commercial immunoassays. We developed an LC-MS/MS based assay for parallel monitoring of multiple peptides in small volumes of human plasma, providing the benefit of analysing exact peptide sequences rather than immuno-reactivity, and potential advantages of cost and sample volumes for measuring multiple peptide hormones. The method involves acetonitrile precipitation of larger proteins, followed by solid phase extraction and nano-LC-MS/MS using an untargeted approach on an orbitrap mass spectrometer. Analysis of plasma from control subjects and patients who have undergone gastrectomy with Roux-en-Y reconstruction, revealed elevated levels of a number of peptides following glucose ingestion. These included GLP-1(7-36), GLP-1(9-36), glicentin, oxyntomodulin, GIP(1-42), GIP(3-42), PYY(1-36), PYY(3-36), neurotensin, insulin and C-peptide, as well as motilin, which decreased following glucose ingestion. Results showed good correlation with those peptides measured previously by immunoassay in the same samples. The gastrectomy group had higher, but non-glucose-dependent, circulating levels of peptides from PIGR and DMBT1.

Overall, the method is fast, generic, reproducible and inexpensive, and requires only small plasma samples, making it potentially adaptable for multiplexed measurement of a variety of peptides.

## Introduction

The gut and pancreas produce a variety of peptide involved in the coordination of intestinal functions, nutrient assimilation, glucose homeostasis and appetite ^1^. Concentrations of peptides in the plasma are altered by fasting and feeding, and are routinely measured for diagnostic and research purposes using immunoassays employing high affinity antibodies. As the gut secretes more than 20 peptides ^2^, research into intestinal physiology is currently hindered by the costs of measuring multiple peptides in parallel and the availability of validated assays. Analysis of individual peptides such as glucagon-like peptide-1 (GLP-1) and PeptideYY (PYY), has revealed that they play substantial roles in the gut-brain and gut-pancreatic axis, and supported the development of GLP-1 based therapies for the treatment of type 2 diabetes and obesity. There is great interest in developing new peptide based therapies for metabolic and intestinal diseases, but gaining a deeper understanding of the physiology of peptide secretion in patients is a critical step in any such drug discovery pathway.

In view of the success of surgical bariatric procedures for the treatment of obesity and type 2 diabetes, there is great interest in understanding the underlying physiological mechanisms. One of the commonest and most effective bariatric procedures is Roux-en-Y gastric bypass (RYGB) surgery, which promotes weight loss (thus increasing insulin sensitivity), and enhances insulin secretion, resulting in rapid resolution of type 2 diabetes with at least partial remission in ~60% of RYGB-patients 1 year post-surgery ^3,4^. The mechanisms underlying these physiological changes are not fully resolved, but a considerable body of evidence points to important roles for peptides such as GLP-I and PYY ^5^, which exhibit profound post-prandial elevations after bariatric surgery. Postsurgical changes in other peptides have been less studied, with some reports of raised and others of unaltered postprandial glucose-dependent insulinotropic polypeptide (GIP) excursions and more sporadic reports on the importance of other peptides which are elevated post-surgically such as Neurotensin (Nts) for surgery outcomes ^7,8^.

The potential for LC-MS based methods to quantify peptides has been demonstrated previously, for example for proglucagon-derived peptides either following immuno-affinity based enrichment ^9^ or after depletion of abundant plasma proteins through solvent precipitation followed by solid phase extraction (SPE)^10^. Whilst these approaches have demonstrated good sensitivity and good correlation with existing immunoassays, they used targeted approaches with triple quadrupole based detection systems, requiring prior knowledge of the analyte under investigation. Alternatively, the plasma peptidome has been investigated after enzymatic protein digestion in an untargeted fashion, but results are at least in part dominated by products from abundant plasma proteins, only partially resolved by specific pre-depletion ^11^ and interpretation might be complicated, when different peptides can be generated from the same prohormone as is the case for proglucagon derived peptides. In our view a better approach for analysing peptides by mass spectrometry in an untargeted fashion is to avoid enzymatic digestion ^12^. We previously reported such an approach for plasma ^13^ from patients with endocrine tumours and for sorted cells ^14^ and tissue extracts from human brain ^15^ and intestines^16^. In this study, we compared changes in the plasma peptidome during an oral glucose tolerance test (OGTT) in subjects after gastrectomy surgery (a procedure resulting in an anatomy very similar to RYGB) and control subjects, using an untargeted LC-MS/MS approach.

## Methods and materials

### Patients and plasma samples

Experiments were performed on plasma samples from a published study, which have previously been analysed by immunoassay for a range of gut and pancreatic peptides ^10^. The study was approved by the National Health Service Research Ethics Committee and conducted in accordance with the ethical standards of the Helsinki Declaration of 1975. In brief, following an overnight fast, all participants drank 50g of glucose in 200ml water within a 5 minute period. Blood samples were collected into EDTA tubes immediately prior to glucose ingestion (time 0), and at 15, 30, 45, 60, 90, 120, 150 and 180 min post-ingestion. Samples were immediately placed on ice and centrifuged for 10 minutes at 3500g at 4°C. 400μl plasma aliquots were snap frozen on dry ice and stored at −80°C within 30 minutes of phlebotomy. Samples from one gastrectomy and one control subject were selected for the pilot study across all timepoints. Samples from 6 gastrectomy and 6 control subjects were selected for the main study analysis at 3 timepoints (0, 30 and 90 min).

### Chemicals

Acetonitrile, methanol, acetic acid and formic acid were purchased from Fisher Scientific, Dithiothreitol, iodoacetamide and bovine insulin were purchased from Sigma Aldrich.

### Extraction of plasma samples

Plasma samples were thawed and extracted on ice to reduce peptidase-based degradation. Plasma (50 μL) was aliquoted in duplicate into a 2 mL 96 well plate and 300 μL of either 80% ACN in water or 80% ACN in water with 0.1% formic acid (FA), both fortified with bovine insulin at 1 ng/mL, were added to each replicate to precipitate high molecular weight proteins; the addition of bovine insulin was only introduced during the main study and omitted in the pilot study. The samples were spun at 3900 g and the supernatants from each time point were combined and transferred to a Lo-bind Eppendorf plate and evaporated under oxygen free nitrogen (OFN) at 40 ⁰C. The dried extracts were reconstituted into 200 μL of 0.1% FA and vortexed before transferring into a Waters Oasis HLB PRiME microelution 96 well plate and washed with 200 μL of 0.1% FA and 200 μL of 5% methanol in 1% acetic acid in water (v/v/v). Peptides were eluted with 2 x 30 μL aliquots of 60% methanol in water with 10% acetic acid (v/v/v). The eluants were dried under OFN and reconstituted with 150 μl of 0.1% FA, of which 40 μl were injected onto the nano-LC-MS system. During the pilot study an additional reduction alkylation was performed after the SPE, which was, however, not found to be necessary for sample analysis and reliable detection of insulin. In brief in the pilot study dried SPE-eluates were dissolved into 75 μL of 50 mM ammonium bicarbonate with 10 mM DTT and reduced at 60 ⁰C for 1 hour. Peptides were alkylated by the addition of 15 μL of 100 mM iodoacetamide in 50 mM ammonium bicarbonate in the dark for 30 minutes. 20 μL of 1% FA was added and 40 μL of sample injected onto the nano LC-MS system.

### Nano LC-MS/MS analysis

Peptide extracts were analysed using a Thermo Fisher Ultimate 3000 nano LC system coupled to a Q Exactive Plus Orbitrap mass spectrometer (Thermo Scientific, San Jose, CA, USA). Extracts were loaded onto a 0.3 × 5 mm peptide trap column (Thermo Scientific) at a flow rate of 30 μL/min and washed for 15 min before switching in line with a 0.075 × 250 mm nano easy column (Thermo Scientific) flowing at 300 nL/min. Both nano and trap column temperatures were set at 45°C. The mobile phases were A: 0.1% FA in water (v/v) and B: 0.1% FA (v/v) in 80:20 ACN/water. Initial conditions were 2.5% B and held for 15 mins. A ramp to 50% B was performed over 90 mins, and the column then washed with 90% B for 20 mins before returning to starting conditions for a further 20 mins, totalling an entire run time of 130 mins. Positive nano electrospray analysis was performed using a spray voltage of 1.8 kV, the tune settings for the mass spectrometer used an S-lens setting of 70 V to target peptides of higher *m/z* values. A full scan range of 400–1600 m/z was performed at a resolution of 75,000 before the top 10 ions of each spectrum were selected for MS/MS analysis. Existing ions selected for fragmentation were added to an exclusion list for 30 s.

### Database searching

LC-MS data were searched using the PEAKS 8.5 software (BSI, Waterloo, Canada) against the human Swissprot database (downloaded 27-10-2017) using a non-specific digest setting. When extracts had been reduced and alkylated, a fixed carboxamidomethylation modification was applied to cysteine residues. Variable modifications included N-terminal acetylation, N-terminal pyroglutamate, C-terminal amidation and methionine oxidation. An FDR setting of 1% was used against a decoy database, and precursor and product ion tolerances were set as 10 ppm and 0.05 *m/z* respectively. The main study cohort data were put through the PEAKSQ software extension to identify potential biomarker peptides in the dataset, manually adding peak areas of bovine insulin as a normalising factor.

### Manual LC-MS data searching and peptide quantitation

The LC-MS/MS and DDA based analysis combined with database searching failed to identify some of the expected gut peptides in all samples. However, in order to obtain a database match, peptides must be both selected for MS/MS fragmentation, and generate a suitably high quality product ion spectrum for the PEAKS software to match against the database. Some gut peptides are present at concentrations in the low pg/ml range, therefore in the presence of other higher concentration plasma peptides, may not be selected for fragmentation. Furthermore, if they were selected for fragmentation, their product ion spectrum might not contain sufficient data for a strong match. Therefore, in order to identify the presence of some of the peptides and to quantify other identified peptides, the theoretical *m/z* values for all peptides were used to interrogate the raw data using the Quan Browser software program (Thermo Scientific) (Table 1). The peak for each peptide was integrated at the expected retention times, with a minimum signal to noise of 3 required, default 9 smoothing added and using the genesis integration algorithm (Example integration shown in figure S1). The data from specific peptides in the large cohort were normalised by expressing their peak areas as a ratio of the internal standard bovine insulin.

**Table 1.** List of peptides selected for manual data searching for quantitation in OGTT samples, along with the *m/z* values associated with the monoisotopic and multiple ^13^C isotope peaks. Peptide *m/z* values were taken from either the PEAKS results or from a previous peptidomics manuscript ^2^. See supplemental data for extracted ion chromatograms, experimentally acquired m/z values and theoretical m/z values for chosen charge states.

| Peptide name | Charge (z) | *m/z* values | RT (mins) |
| --- | --- | --- | --- |
| GRPP | 4 | 846.63, 846.88, 847.13, 847.38 | 43 |
| OXN | 7 | 636.46, 636.60, 636.75, 636.89 | 58 |
| Glicentin | 7 | 1157.84, 1157.98, 1158.13, 1158.27, 1158.41, 1158.56 | 56 |
| GLP-I 7-36 amide | 4 | 824.91, 825.18, 825.43 | 72 |
| GLP-I 9-36 amide | 4 | 772.90, 773.15, 773.40, 773.65, 773.90 | 77 |
| PYY 1-36 amide | 7 | 616.60, 616.75, 616.89, 617.04 | 60 |
| PYY 3-36 amide | 7 | 579.30, 579.45, 579.59 | 58 |
| Insulin (intact) | 5 | 1162.14, 1162.34, 1162.54, 1162.74, 1162.94 | 66 |
| Insulin A-chain | 2 | 1306.05, 1306.55, 1307.05, 1307.55 | 66 |
| Insulin B-chain | 5 | 709.35, 709.55, 709.75, 709.95, 710.15 | 60 |
| C-peptide | 3 | 1007.17, 1007.51, 1007.84, 1008.18 | 75 |
| Bovine insulin (intact) | 5 | 1147.13, 1147.33, 1147.53, 1147.73, 1147.93, 1148.13 | 65 |
| GIP propeptide | 7 | 455.24, 455.39, 455.53, 455.67 | 31 |
| GIP 1-42 | 6 | 831.25, 831.41, 831.58, 831.75, 831.91 | 62 |
| GIP 3-42 | 7 | 792.23, 792.40, 792.56, 792.73, 792.90 | 60 |
| Neurotensin | 3 | 558.31, 558.64, 558.98 | 43 |
| Motilin | 6 | 540.48, 540.68, 540.88, 541.08 | 48 |
| Adrenomedullin (45-92) | 8 | 640.08, 640.20, 640.45, 640.58 | 48 |
| Augurin | 6 | 498.11, 498.28, 498.45, 498.61 | 42 |

### Immunoassays

Immunoassays on these samples have been described previously ^10^. In brief, total GLP-1, total GIP and total PYY were measured using Mesoscale Discovery (MesoScale Diagnostics, Rockville, USA) assays according to the manufacturer’s instructions. Insulin concentrations were measured using the Diasorin Liaison XL insulin system (Diasorin, Milan, Italy).

## Results

### Full OGTT sample comparison in two subjects

A pilot study was performed to examine the plasma peptidome at multiple time points following a 50g OGTT in one gastrectomy subject and one healthy control subject, generating data about which peptides were detectable, which appeared different following gastrectomy, and the best time points for further testing in a main study cohort. A number of circulating peptides were detected, including gut and pancreatic peptides, fibrinogen fragments, hepcidin, thymosin, bradykinin and angiotensin. Many of the identified gut derived peptides showed clear time dependent changes after on OGTT, as shown in Figure 1. Previously published ELISA results from these 2 subjects are shown in Figure S2.

**Figure 1.**
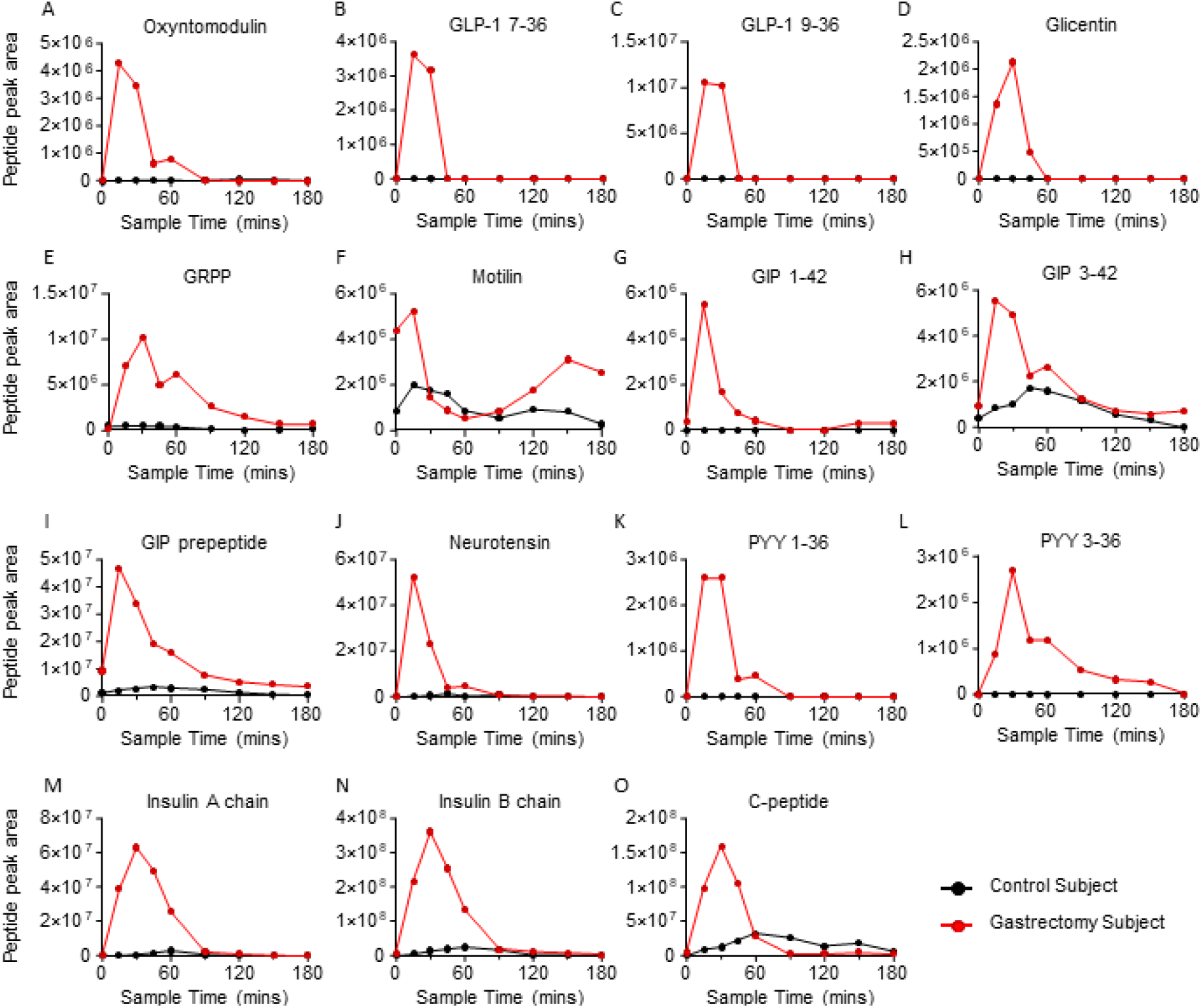
Plasma peptide peak area time courses for one gastrectomy and one control participant in the pilot study. Plasma concentrations of the peptides indicated, measured by LC-MS/MS and quantified as peak areas, in the 2 subjects at the indicated timepoints after a 50g OGTT.

Extracts from the gastrectomy subject returned more gut peptide identifications than the control subject, in particular peptides from the proglucagon gene, which included OXN, active GLP-1 7-36 amide and GLP-1 9-36 amide, the inactive cleavage product of dipeptidyl peptidase-4 (DPP4) digestion, their levels over the OGTT can be seen in Figures 1A-C. Another intestinally proglucagon-derived peptide, glicentin, comprising the N-terminal part of proglucagon, including both the GRPP and oxyntomodulin sequence, which has 68 amino acids - too large to be identified by PEAKS, which has a cut-off of 65 amino acids. Manual examination of the raw data for the expected *m/z* of its [M+7H]^7+^ charge state showed glicentin was present in the gastrectomy subject, but not the control subject (Figure 1D). The only other proglucagon derived peptide we were able to detect was GRPP, which was detectable in both the post-gastrectomy subject and the control subject, if at lower levels in the latter (Figure 1E).

The analysis also identified peptides from GIP and motilin, a peptide involved in gut motility ^17^, both of which are expressed at highest levels in the proximal small intestine (duodenum/jejunum) ^2^. Motilin was readily detectable in both subjects and levels dropped after glucose ingestion in both cases (Figure 1F). With the exception of active GIP 1-42 (Figure 1G), which we could not detect in the control subject, GIP-derived peptides (GIP-prepeptide, representing the first cleaved peptide, N-terminal to the active hormone and GIP 3-42, an inactive product of GIP 1-42 digestion by DPP4) (Figure 1H,I) were readily detectable in both subjects and rose after the OGTT, reaching higher levels in the gastrectomy samples. Neurotensin, a peptide thought to arise mostly from the distal small intestine, rose after the OGTT in both subjects, but to a greater extent and earlier in the gastrectomy subject (Figure 1J). PYY, a peptide expressed at higher levels in the more distal intestine was only identifiable in the plasma database search after correction of the Uniprot database. Whereas the first 8 N-terminal amino acids of PYY 1-36 are designated as IKPEAP**R**E in the standard Uniprot database, the correct sequence is IKPEAP**G**E. Both full length PYY 1-36 amide and DPP4-cleaved PYY 3-36 amide (Figure 1K,L) were detected in the gastrectomy, but not the control subject.

The LC-MS analysis readily detected insulin A, B and C peptides (Figure 1 M-O), the time-profiles of which mirrored the immunoassay derived insulin concentrations in the same samples published previously (Figure S2, ^10^). Plasma concentrations of insulin and its C-peptide are about 2 orders of magnitude higher than those of proglucagon-derived peptides, and are well within the sensitivity of the LC-MS system, as we have previously demonstrated ^18^. A correlation of the insulin ELISA concentration against the B-chain peptide peak area across these 2 participants gave an R^2^ value of 0.9849, showing very comparable data between the two analytical approaches, despite the LC-MS data not being transformed or normalised in any way.

### Peptidomics analysis of 6 gastrectomy and 6 control subjects

We next analysed plasma samples from 6 gastrectomy subjects and 6 control cubjects taken at 0, 30 and 90 min after the 50g OGTT, using the same analytical protocol but omitting the reduction/alkylation step (leaving insulin and other disulphide bonded peptides intact). The PEAKS output identified similar peptides to those found in the pilot study, including a number of distinct products from the PYY, proglucagon and GIP genes, together with motilin and C-peptide. Peak areas of all previously described peptides were subsequently generated by interrogation of the raw data using the Quan Browser software, along with peak areas of intact insulin and the internal standard bovine insulin. The addition of bovine insulin as an internal standard enabled normalisation of the peak area for each peptide to that of bovine insulin in the same sample, generating semi-quantitative peak area ratios for each analyte. The peak areas of the detected bovine insulin internal standard over the 36 nano LC-MS analyses was consistent, with a %CV of 11.9%, indicating that the extraction process and LC-MS analysis were robust and reproducible.

Peak area ratios of the selected peptides at 0, 30 and 90 min after the OGTT are depicted in Figure 2. Confirming the pilot data and previously generated results from immunoassays ^10^, plasma levels of insulin, PYY, proglucagon-derived peptides and neurotensin were elevated after the OGTT in gastrectomy subjects compared with control subjects, whereas GIP-derived peptides were largely similar between the groups. As we have previously analysed several gut hormones in these samples by immunoassays, using commercial kits against total GLP-1 (GLP-I 7-36 and GLP-I 9-36 combined), total GIP (GIP 1-42 and GIP 3-42 combined), total PYY (PYY 1-36 and PYY 3-36 combined) and insulin, we examined the performance of the LC-MS approach, by comparing LC-MS results against the corresponding immunoassay values (Figure 3). Across the 12 subjects, there was excellent correlation between insulin levels measured with the two assay methods (R^2^ = 0.9837). Immunoassay-determined total GLP-1 correlated reasonably with the peak area ratio for GLP-1 9-36 amide (the major circulating form of GLP-1 detected by LC-MS; R^2^ = 0.73), and slightly better with the sum of GLP-1 7-36 and GLP-1 9-36 (R^2^ = 0.80) or GRPP (R^2^ = 0.84), which is co-released from intestinal L-cells but can also arise from glucagon-producing pancreatic alpha-cells. Individual GIP-derived peptide LC-MS data were correlated with total GIP immunoassay results, but also improved when summing results from both the GIP 1-42 and 3-42 peptides. GIP prepeptide, comprising the cleaved peptide cleaved from the prohormone on the N-terminal side of GIP, was more readily detectable by LC-MS than GIP 1-42 or 3-42 but its levels correlated less well with the immunoassay than GIP 3-42 or total GIP. Total PYY measured by immunoassay correlated well with peak area ratios for PYY 3-36 amide (R^2^ = 0.93) and total PYY (1-36 amide + 3-36 amide; R^2^ = 0.94), whereas the correlation against PYY 1-36 amide alone was weaker at R^2^ = 0.78.

**Figure 2.**
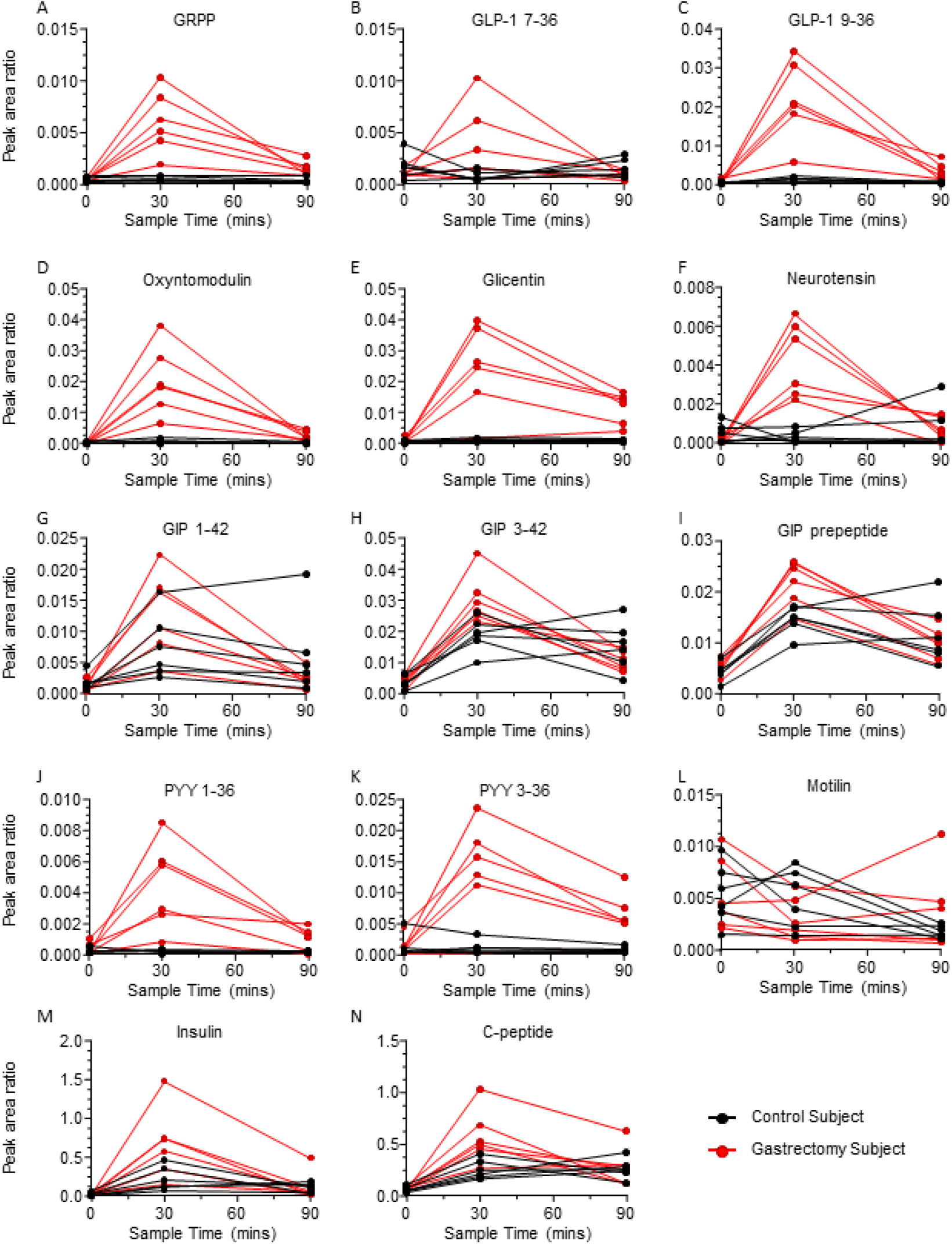
Peptide peak area ratios from a group of control and gastrectomised individuals after OGTT. Peak area ratios for the peptides indicated, measured by LC-MS/MS. Samples were taken at t=0, 30 and 90 min following a 50g OGTT in 6 healthy control and 6 gastrectomised individuals.

**Figure 3.**
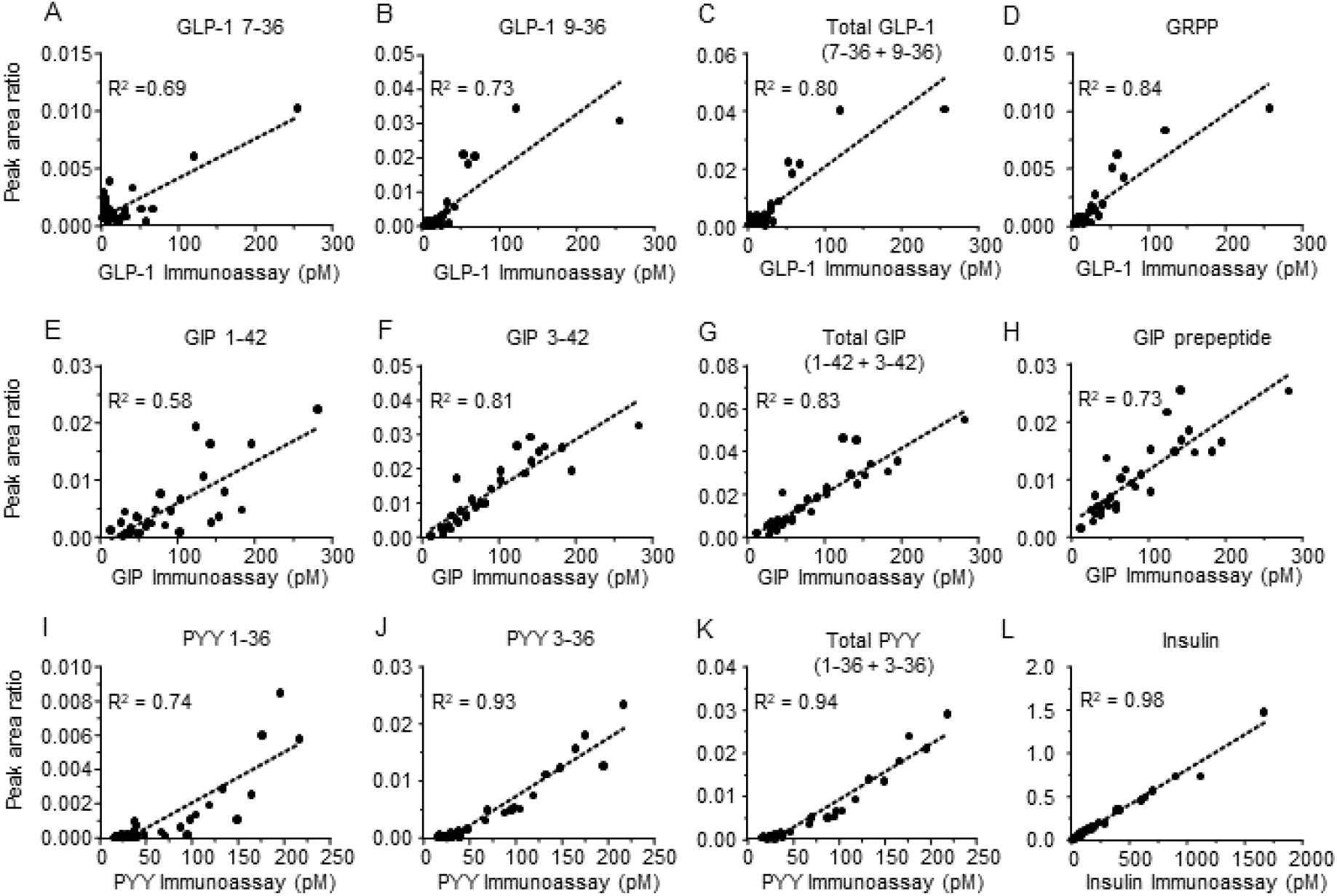
Correlations between plasma peptide measurements by LC-MS/MS and immunoassay. Data from the 0, 30 and 90 min timepoints after the OGTT in control and gastrectomy subjects from Ffigure 2 are plotted as peak area ratios (y-axis) for the peptides indicated against previously-measured immunoassay levels in the same samples ^10^. The immunoassays used were the total GLP-1, total GIP and total PYY assays from MesoScale Discovery. Dashed lines show the line of best correlation, and correlation coefficients (R^2^) are shown for each plot.

### PEAKSQ analysis from 6 gastrectomy and 6 control subjects

Analysis using the PEAKSQ module, manually adding the bovine insulin peak area as a normalising factor, enabled interrogation of the entire dataset for plasma peptidomic differences between control and gastrectomy subjects. Across all timepoints, a number of likely false positives were detected, including peptides from fibrinogen and globins, and closer interrogation of the data revealed that the PEAKSQ software was selecting up to three peptides for quantitation out of the many identified from the same parent protein, whereas other peptides from the same parent exhibited either no differences or changes in the opposite direction. In the dataset collected 30 min after glucose ingestion, PEAKSQ correctly identified that insulin, GIP and proglucagon products were higher in gastrectomy than control participants, supporting the data presented in Figures 1 and 2, but did not identify differences in any other known and identified bioactive peptides. Across all samples, PEAKSQ identified raised levels of peptide fragments from two larger proteins, PIGR (polymeric immunoglobulin receptor) and DMBT1 (Deleted in malignant brain tumours 1) (Figure 4), which are both known to be enriched in the gastrointestinal tract ^19^. PIGR has a large extracellular domain involved in binding IgA and IgM for translocation across epithelia, a single transmembrane domain and a short intracellular C-terminus. Peptides identified from PIGR included the sequences 598-639, 598-648, 604-639, 605-648, 607-648, 610-648 and 604-648, which mainly comprise of the C-terminus of the extracellular domain (19-638) and part of the transmembrane domain (639-661). Across all time points, all peptides derived from PIGR had higher peak areas in gastrectomy than control subjects (e.g. p=0.009 for fragment 598-648 by 2-way ANOVA). One peptide fragment from DMBT1 (amino acids 2385-2413 from the C-terminus) was higher in plasma from gastrectomy patients than controls (p=0.017 by 2-way ANOVA). Figure 4 also shows the peak areas of two unrelated peptides that were detected in the plasma for comparison. For these two peptides, Augurin 42-68 (a C-terminal fragment of a propeptide containing residues 32-68) and Adrenomedullin 45-92 which is an intact propeptide sequence, where no significant differences between the control and gastrectomised subjects were observed.

**Figure 4.**
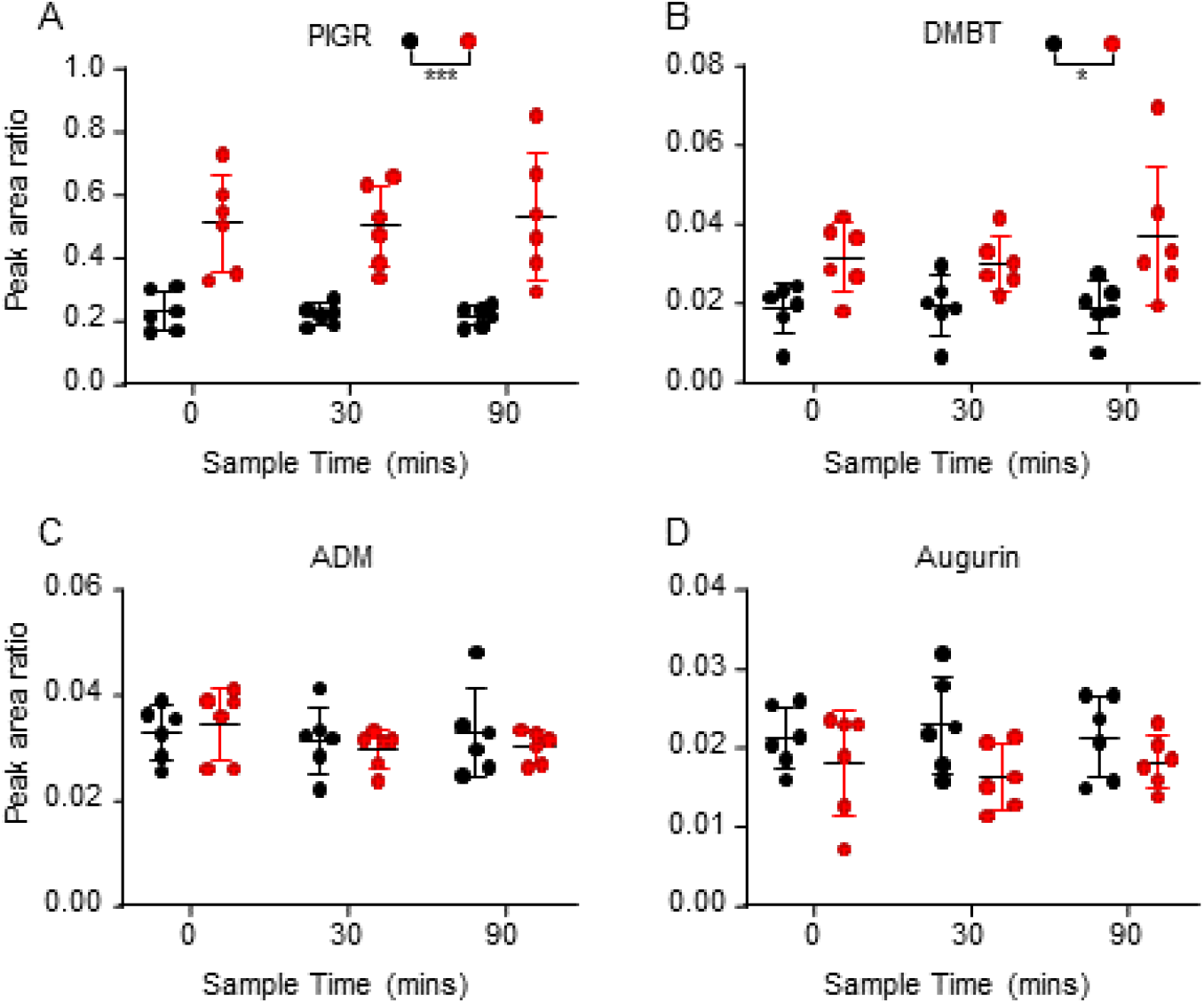
Peak area ratios of plasma peptides that were glucose independent. Peak area ratios for the peptides indicated, measured by LC-MS/MS. Samples were taken at t=0, 30 and 90 min following a 50g OGTT in 6 healthy control and 6 gastrectomised individuals. PIGR 598-648 is illustrative of a range of PIGR peptides, and together with DMBT 2385-2413 was higher in gastrectomy than control individuals. Augurin (42-68) and ADM (45-92) are shown as examples of peptides that did not differ between groups.

## Discussion

We have demonstrated that a simple, high throughput and inexpensive plasma peptide extraction methodology can be combined with nano LC-MS/MS analysis to study gross changes in the plasma peptidome following gastrectomy surgery. Insulin and C-peptide were readily detectable by LC-MS/MS in all participants and correlated well with immunoassay derived concentrations across the full range of the fasting and stimulated levels found in the subjects tested. Gut peptides circulate at lower concentrations than insulin, and were more difficult to detect by LC-MS/MS, although in the gastrectomy samples we readily identified a number of gut hormones believed to contribute to metabolic improvements after bariatric surgery (GLP-I, OXN, GIP, PYY and Neurotensin). We previously reported substantially elevated plasma levels during an OGTT in this group for GLP-1 and PYY, measured by immunoassays and the multiplex LC-MS/MS results reported here correlate well with the historical data. Post-gastrectomy anatomy is strikingly similar to post-RYGB anatomy (with the additional total removal of the stomach remnant in the former), but in the predominantly lean post-gastrectomy cohort the excessive insulin secretion often results in postprandial hypoglycaemia, which can be alleviated by GLP1R-blockage ^16^, with similar observations having been reported in RYGB-patients after weight-loss ^20^.

Most gut peptides, however, were below the detection limit in control subjects, and only identifiable in a few control subject samples after interrogation of the raw data. For example, the active form of GLP-1 (GLP-1 7-36 amide) was not detected in plasma from the control subject as normal concentrations are less than 10 pmol/l ^21^, below the detection limit of our generic nano LC-MS approach. However, where immunoassay data were available, the LC-MS peak area ratios for post-OGTT gut hormone levels in the gastrectomy group correlated well with immunoassay-derived concentrations. This indicates that the developed method is inherently quantitative, and is therefore applicable for the further development of fully quantitative studies using peptide standards and stable isotope labelled internal standards ^10^. Arguably, our results suggest that nano LC-MS approaches are well suited for monitoring multiple peptides in parallel in small plasma samples. The sensitivity achieved was in the low tens of picograms per millilitre of plasma, which could potentially be improved upon by using more targeted SRM based analysis rather than the full scan function used in the data dependent acquisition technique on the Orbitrap. Indeed, we have reported post-OGTT glucagon excursions quantified by LC-MS/MS in these patients and the controls using a targeted approach ^10^. However, in its current guise, the approach could be used for monitoring gut peptide release in post-gastric bypass patients. Other groups have studied the plasma peptidome of patients post RYGB by LC-MS and detected raised oxyntomodulin (OXN) levels ^11^, but their approach involved substantial sample work-up including enzymatic digestion of target peptides which complicates interpretation of the source of the resulting peptides as OXN also contains sequences present in both glucagon and glicentin (two other peptides produced from the proglucagon gene, which are easily distinguished by our method). In this context, although the presence of raised plasma glucagon after gastric bypass has been reported ^22^, our analysis did not identify glucagon in the plasma despite elevated glicentin and OXN (Figure 1) and we previously have shown similar relatively small glucagon-excursions in the control and post-gastrectomy patients ^10^.

The peptidomics approach identified some non-glucose dependent peptides that appeared to differentiate the gastrectomy from the control subjects. These peptides were derived from proteins that have been attributed to host responses to infection in the gut (PIGR and DMBT1) and were higher in concentration in the gastrectomy group. PIGR binds polymeric immunoglobulins on the basolateral epithelial surface, transports them to the apical membrane in vesicular structures, and is then cleaved to release its immunoglobulin cargo into the gut lumen ^23^. Our detection of PIGR-derived peptides in the plasma likely reflects either cleavage occurring at the basolateral membrane, or reabsorption of cleaved PIGR peptides from the lumen. Our finding that several circulating PIGR-derived peptides contained part of the transmembrane domain suggests that cleavage may involve gamma-secretase. Release of the PIGR extracellular domain or PIGR-derived peptides into the circulation has not been described previously, and further studies will be required to determine whether PIGR-derived plasma peptides have biological activity. DMBT1 is large secreted protein found in intestine and saliva ^24^ and is also known as salivary agglutinin, surfactant pulmonary-associated D-binding protein, and Hensin. DMBT1 is reported to play a role in intestinal microbial defence ^25^, and is upregulated in individuals with irritable bowel disease and other diseases of the intestinal tract ^26^. As both PIGR and DMBT1 have been implicated in host defence, we speculate that their detection at higher levels post gastrectomy may reflect an increased load of intestinal microbiota, as gastrectomy patients frequently suffer from small intestinal bacterial overgrowth and PIGR expression is known to be activated by microbial products such as LPS ^27^.

## Conclusions

The described peptidomics approach employs a generic protein precipitation approach (followed by SPE) for enriching for circulating peptides whilst removing high abundance and high molecular weight proteins such as albumin and immunoglobulins. The approach requires only small volumes of plasma (50 μl) and is a fast, generic, reproducible and inexpensive method for studying an under-researched area, the plasma peptidome. Whilst the method was used in a semi quantitative fashion, it generated data that showed a good correlation with existing immunologically derived plasma peptide concentrations. The peptidomics analysis identified similar increases in known bioactive peptides after gastrectomy, however the sensitivity of the approach wasn’t sufficient to detect peptides circulating in the low pg/ml concentrations in the control subjects. Whilst no new glucose dependent peptide changes were detected, peptides from DMBT and PIGR appeared raised after gastrectomy, and will require further studies to investigate whether they could be used as biomarkers for intestinal infection or inflammation.

## SUPPORTING INFORMATION

Figure S1. Typical integration of insulin in control subject time point of 30 minutes post glucose ingestion. Image shows peak retention time and MS spectra of intact insulin.

Figure S2. Immunoassay derived plasma concentrations of insulin, GIP (total measurement of 1-42 and 3-42), GLP-I (total measurement of 7-36 and 9-36) and Peptide YY.

## Acknowledgements

We thank the participating patients and the Cambridge Oesophago-gastric unit, the NIHR Clinical Research Facility (CRF) and the Translation Research Facility (TRF) and Keith Burling, Peter Barker and the staff at the NIHR BRC Core Biochemical Assay Laboratory (supported by the MRC [MRC_MC_UU_ 344 12012/5] and Wellcome Trust [100574/Z/12/Z]) for support of this study.

---

**Supplemental Figure S1.**
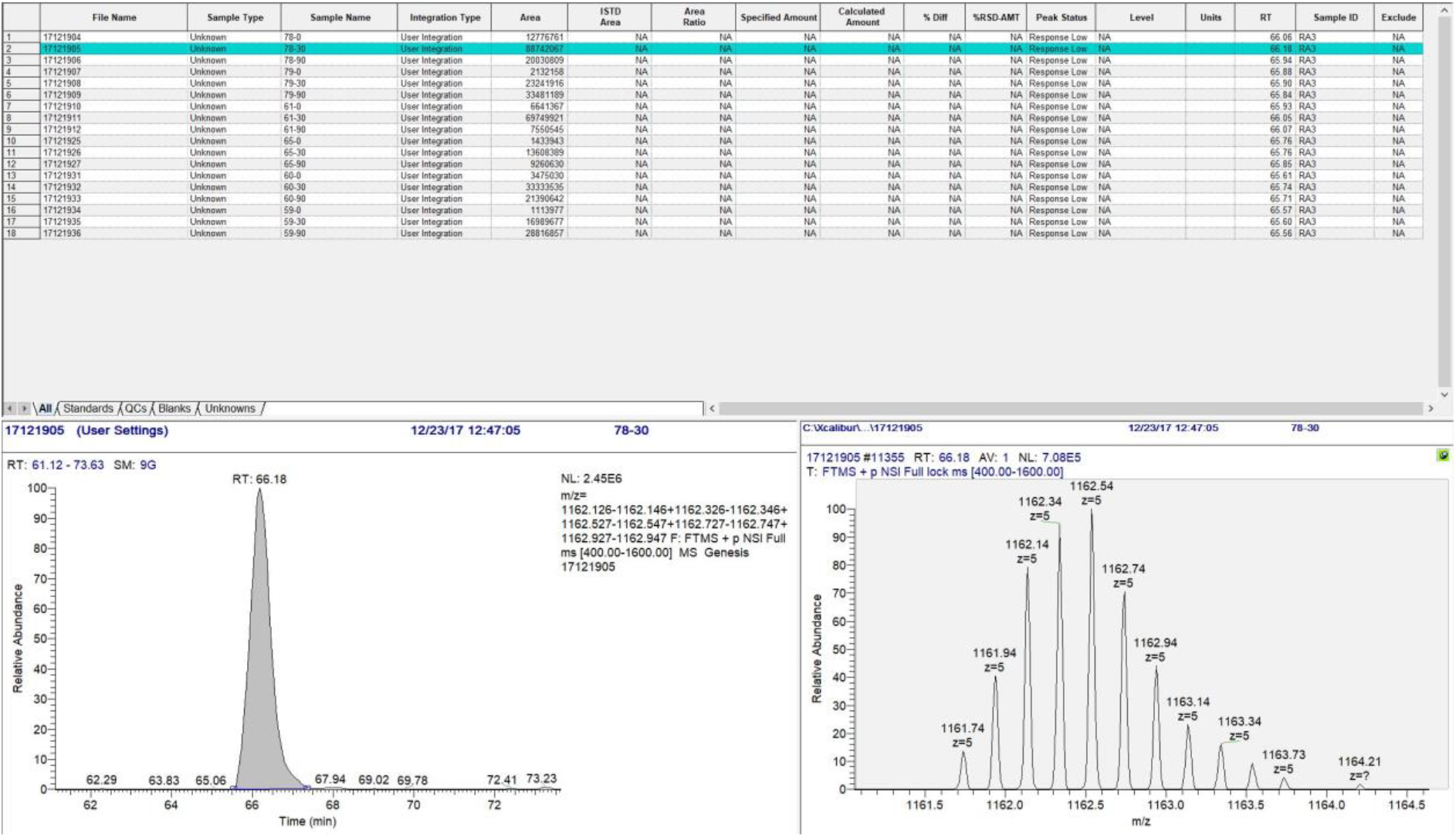
Example integration of insulin peptide in a 30 minute time point sample from a control subject. Figure shows intact insulin peak, corresponding peptide isotope pattern and peak areas associated with this peptide in control subjects.

**Supplemental Figure S2.**
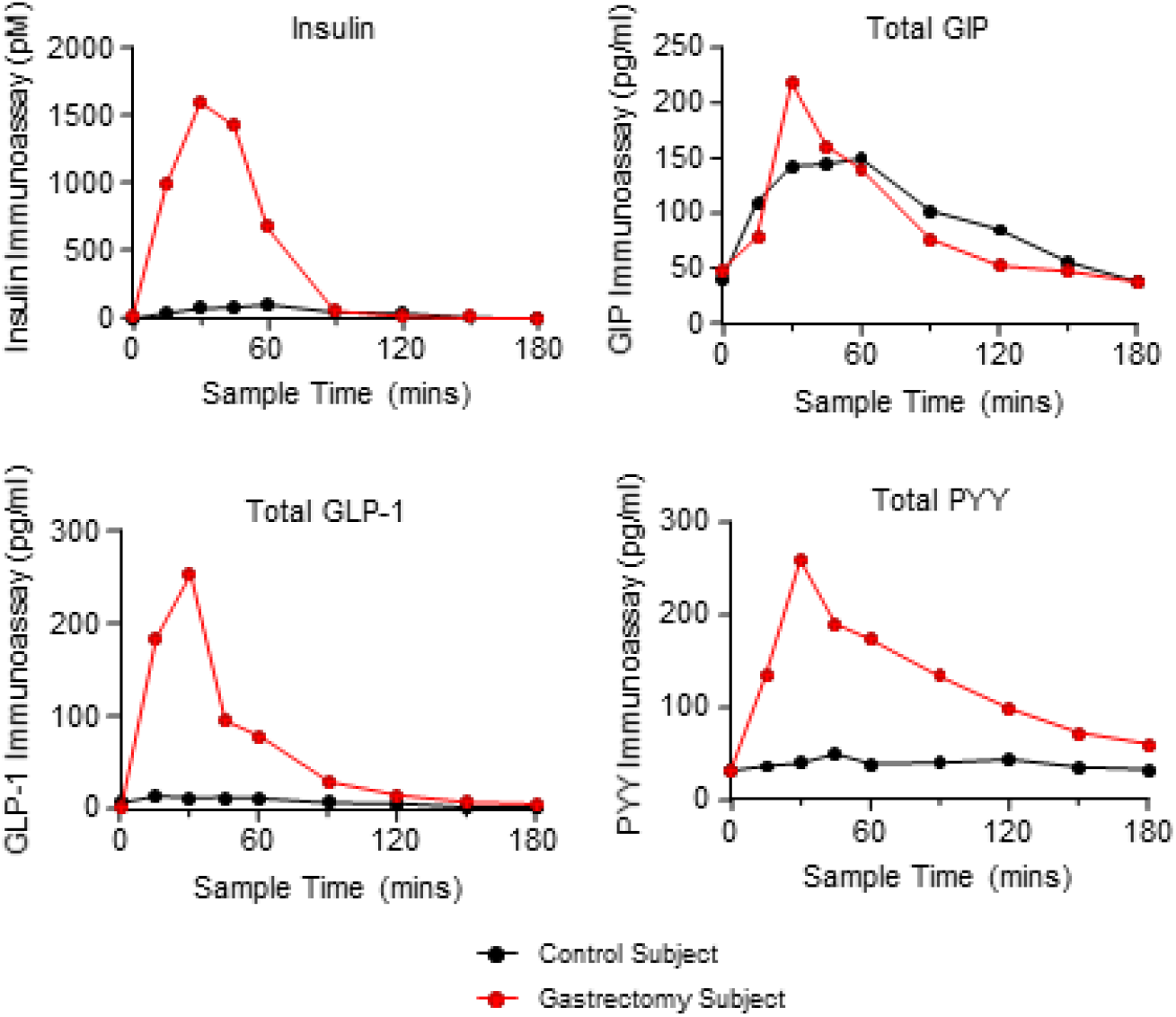
Immunoassay derived peptide concentrations over an OGTT. Plasma concentrations of insulin, GIP (total measurement of 1-42 and 3-42), GLP-I (total measurement of 7-36 and 9-36) and Peptide YY. Data clearly shows how gastric bypass results in increased release of insulin, GLP-I and PYY peptides in response to a glucose challenge.

