## supplemental data for "Mass spectrometric characterisation of the circulating peptidome following oral glucose ingestion in control and gastrectomised patients"

### GRPP

C:\Xcalibur\...\17121938 *2/26/17 12:47:54 69-30

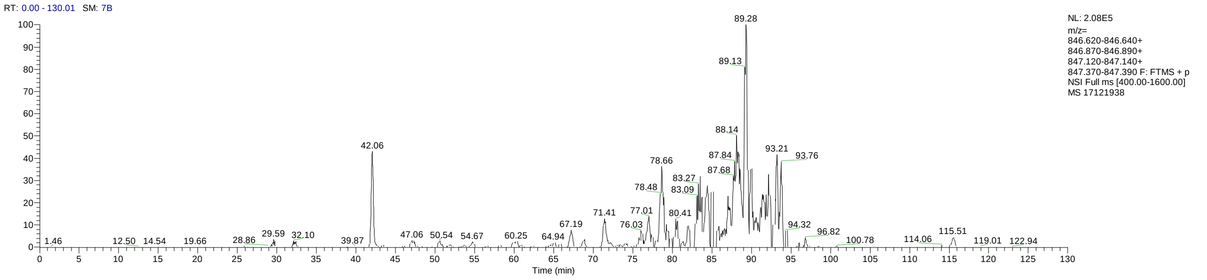


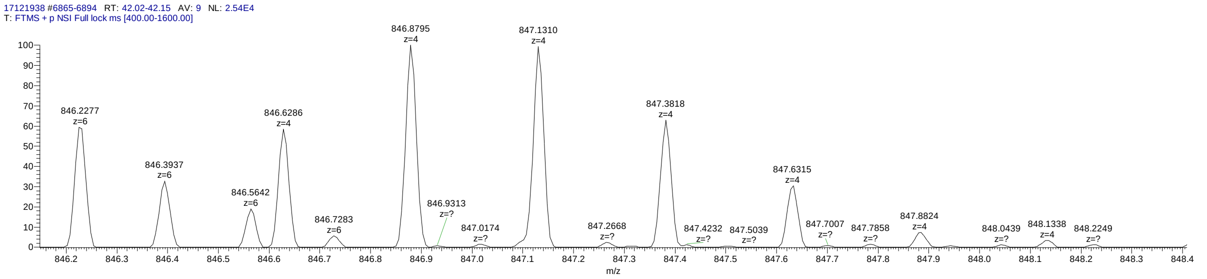


RSLQDTEEKSRSFSASQADPLSDPDQMNED +H +H2O: C136 H219 N41 ...

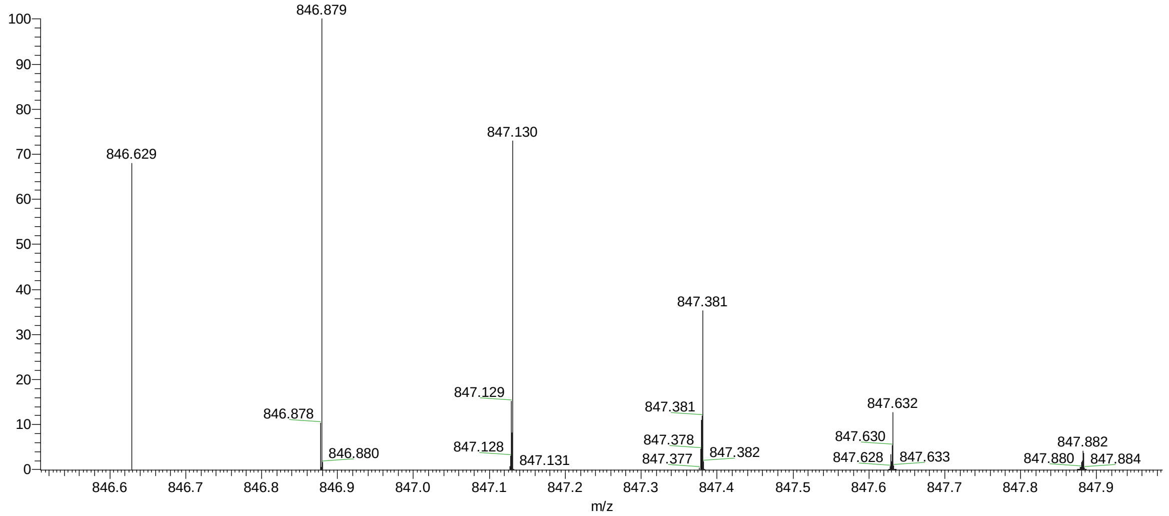

### Oxyntomodulin

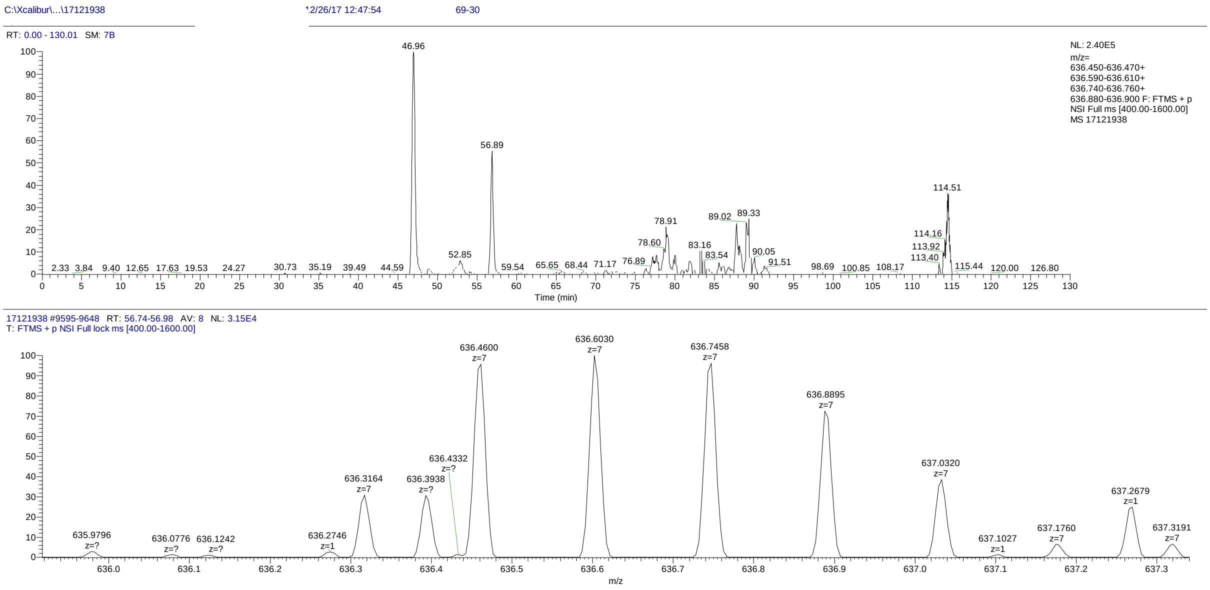


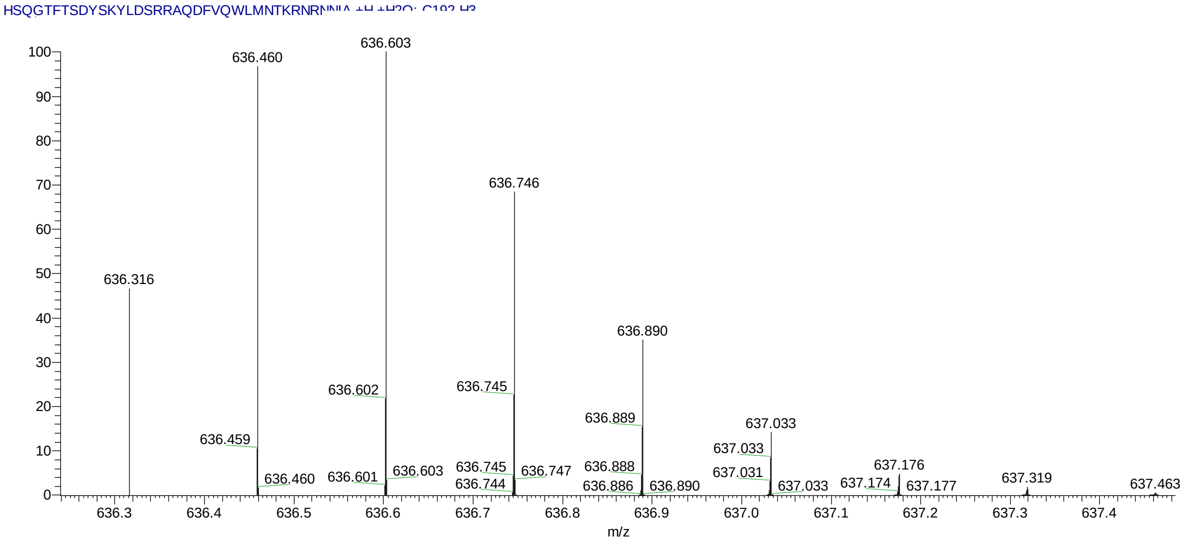

### Glicentin

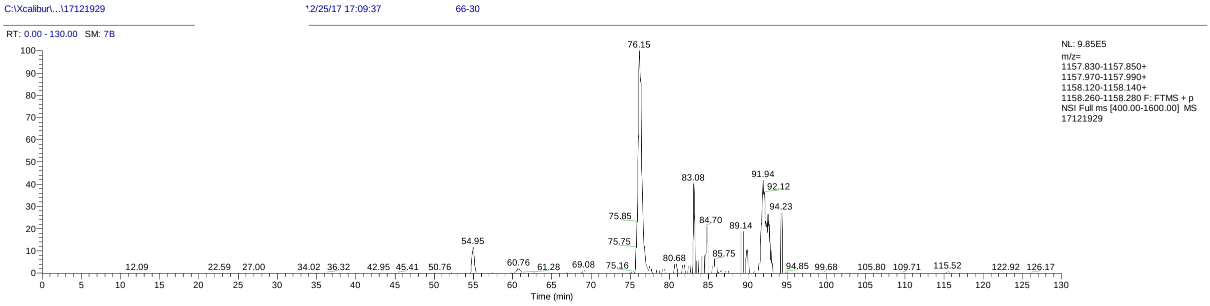


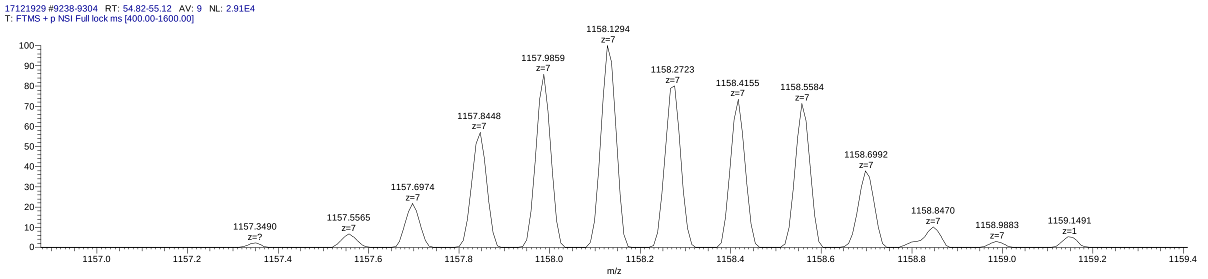


### Theoretical isotope pattern for $[M+7H]^{7+}$ charge state

RSLQDTEEKSRSFSASQADPLSDPDQMNEDKRHSQGTFTSDYSKYLDSRRAQDFVQ...

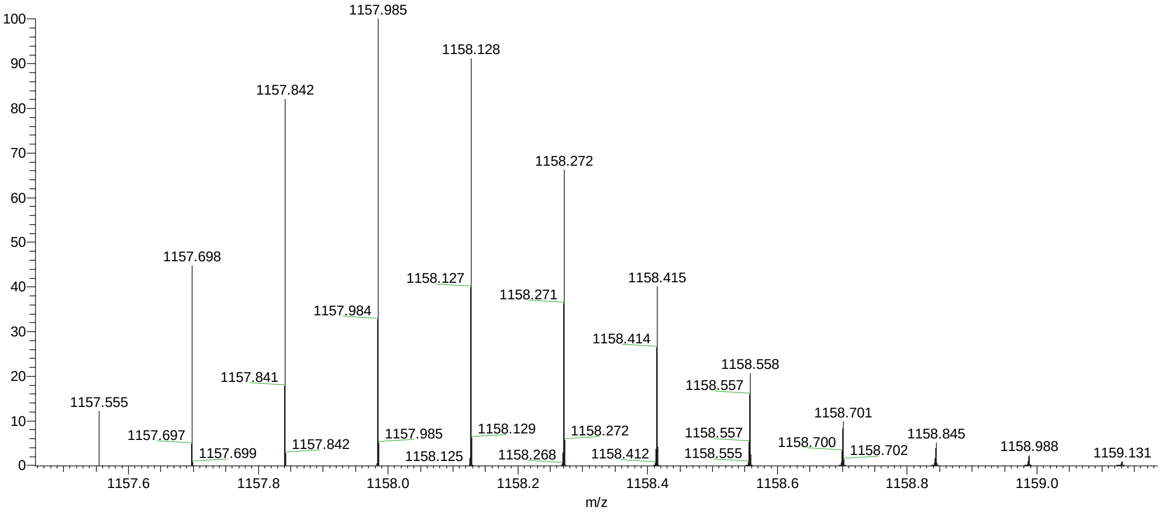

### GLP-I 7-36 amide

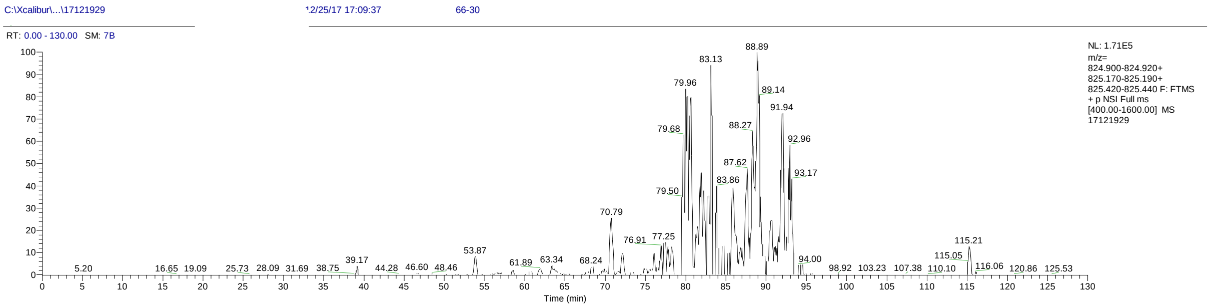


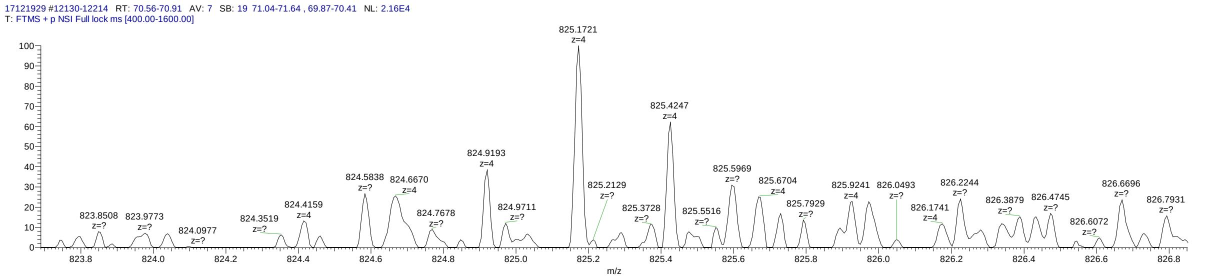


#### Theoretical isotope pattern for $[M+4H]^{4+}$ charge state

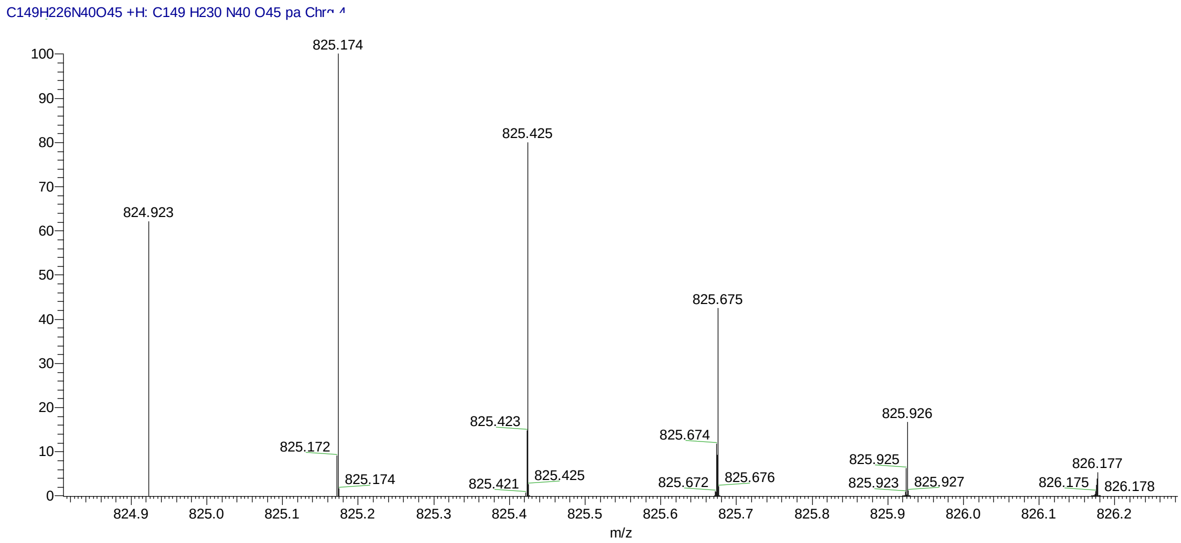

### GLP-I 9-36 amide

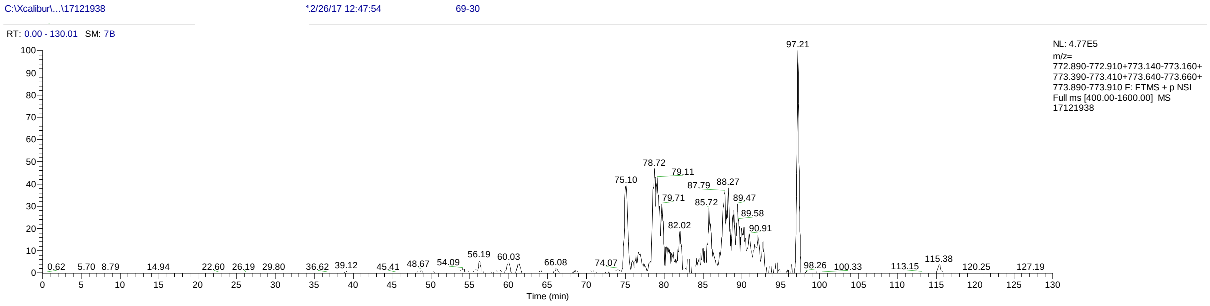


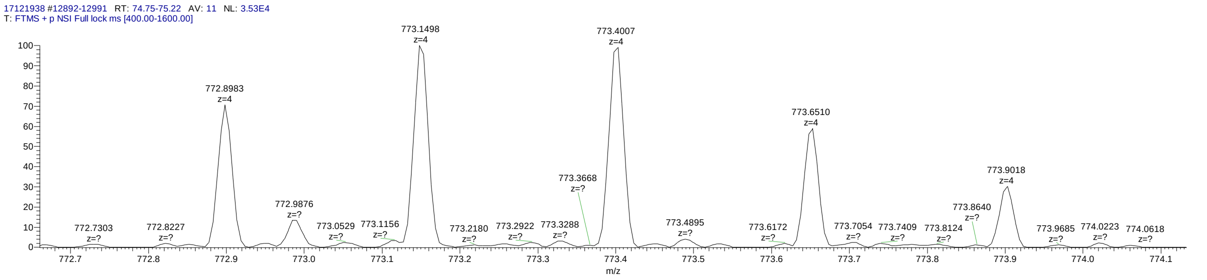


### Theoretical isotope pattern for $[M+4H]^{4+}$ charge state

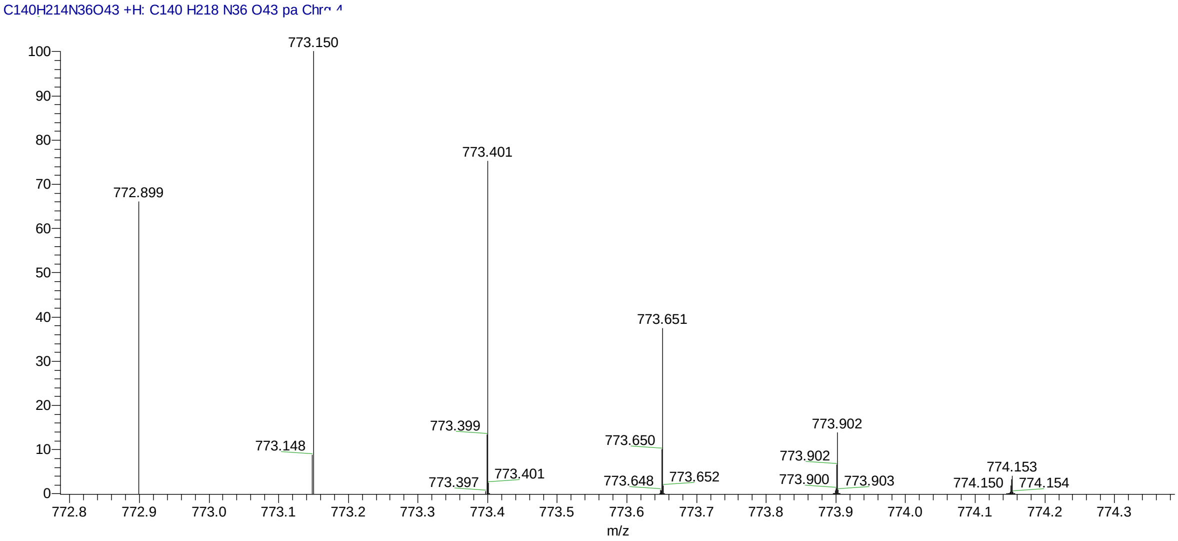

### PYY 1-36 amide

#### YPIKPEAP**G**EDASPEELNRYYASLRHYLNLVTRQRY

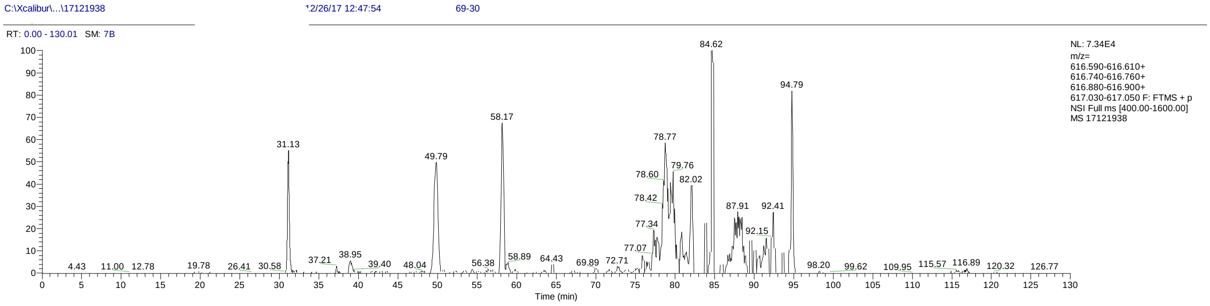


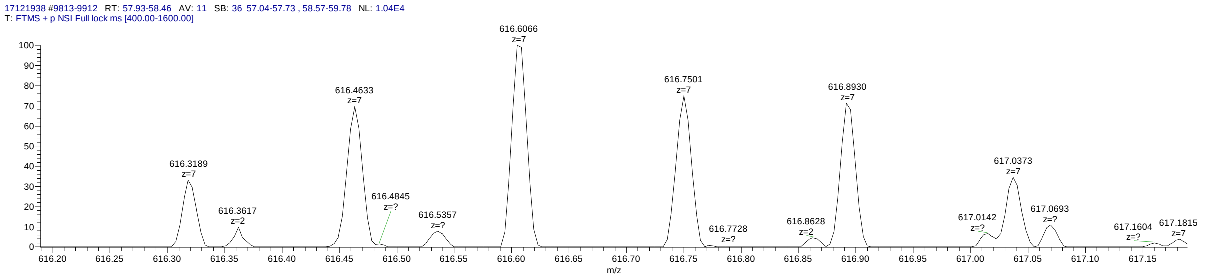


#### Theoretical isotope pattern for $[M+7H]^{7+}$ charge state

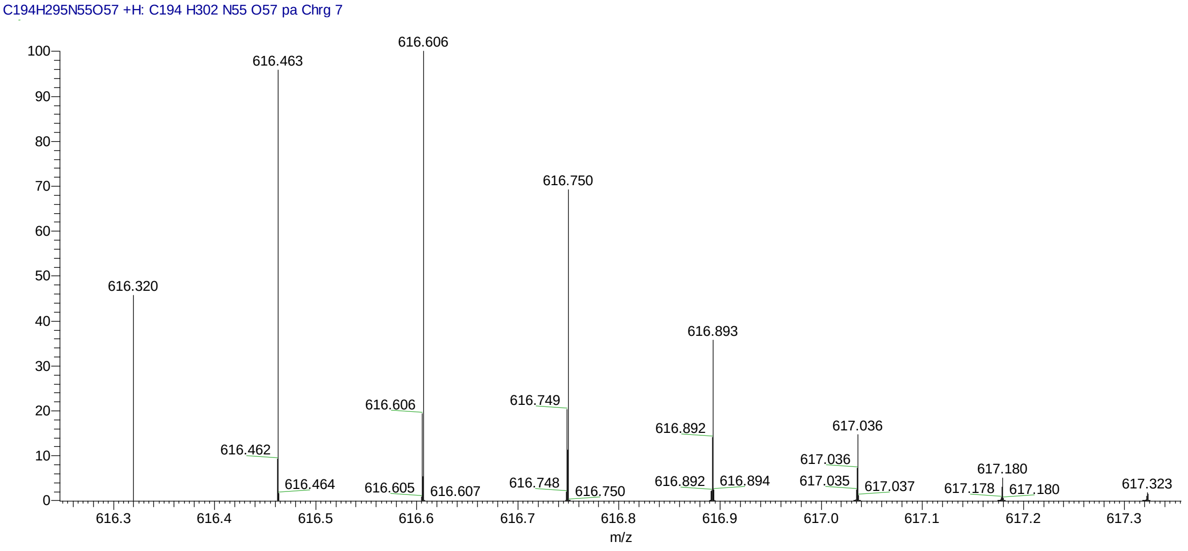

### PYY 3-36 amide

#### IKPEAP**<u>G</u>**EDASPEELNRYYASLRHYLNLVTRQRY

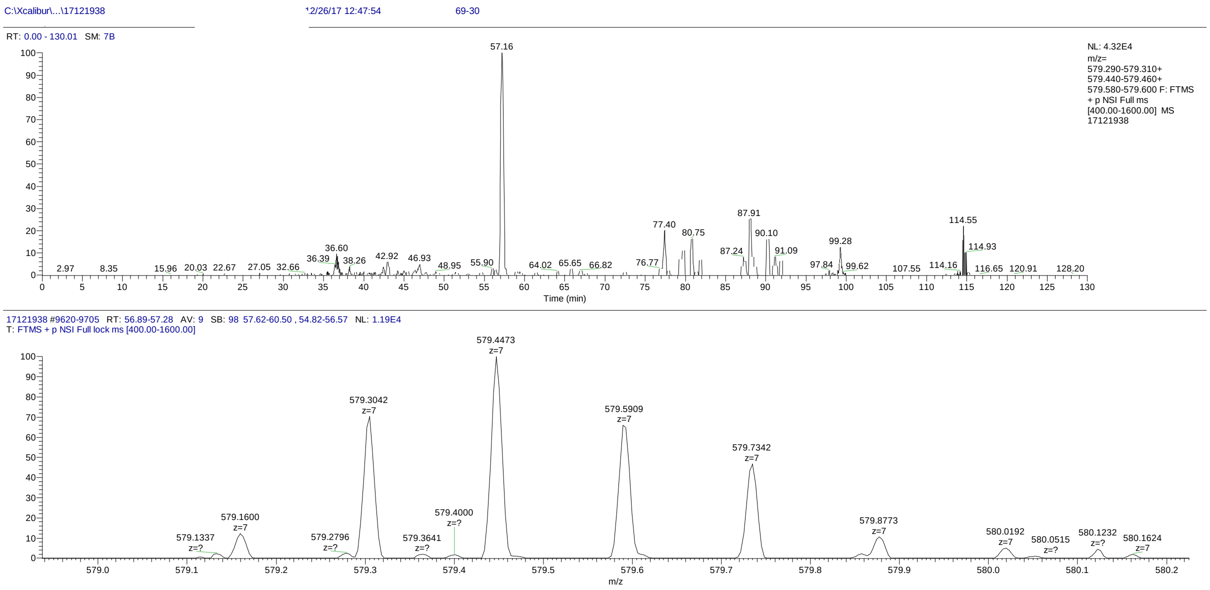


#### Theoretical isotope pattern for $[M+7H]^{7+}$ charge state

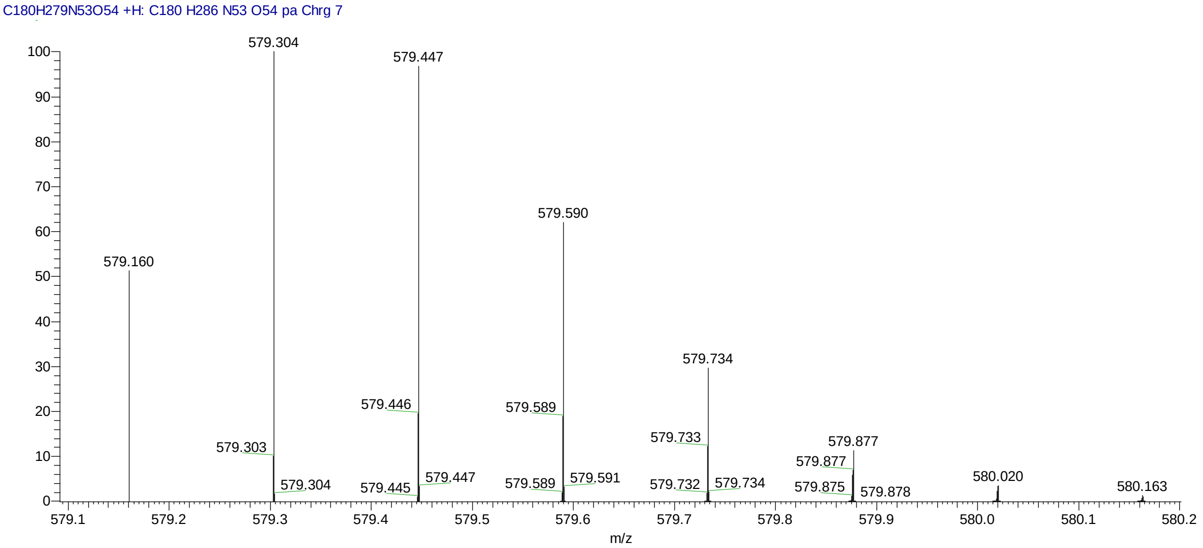

### Intact human insulin

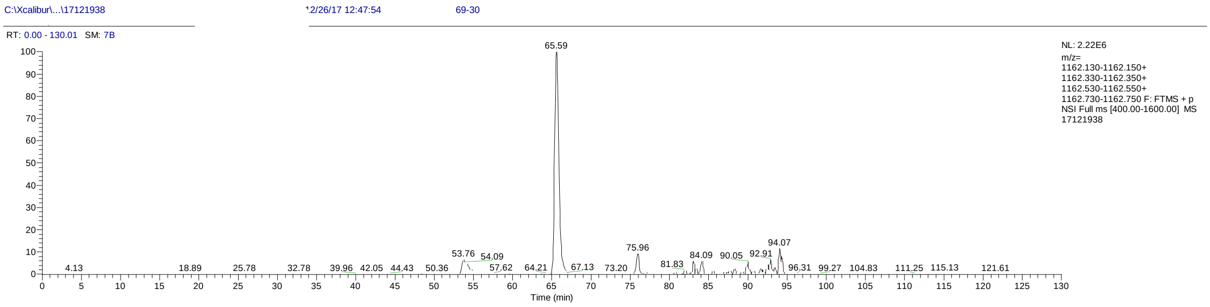


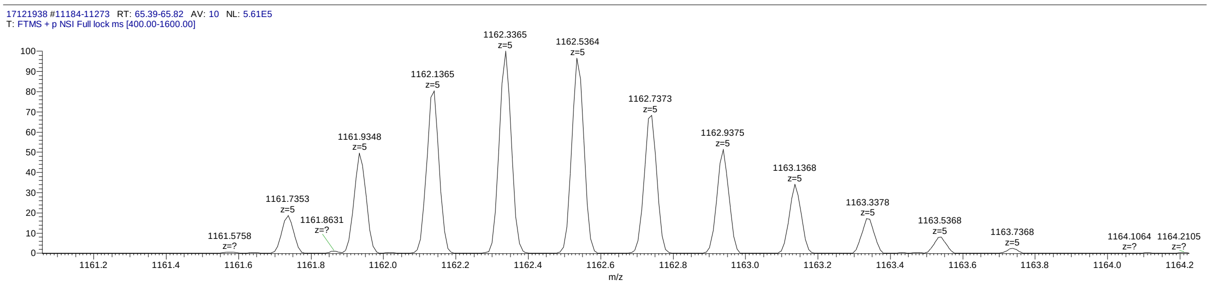


### Theoretical isotope pattern for $[M+5H]^{5+}$ charge state

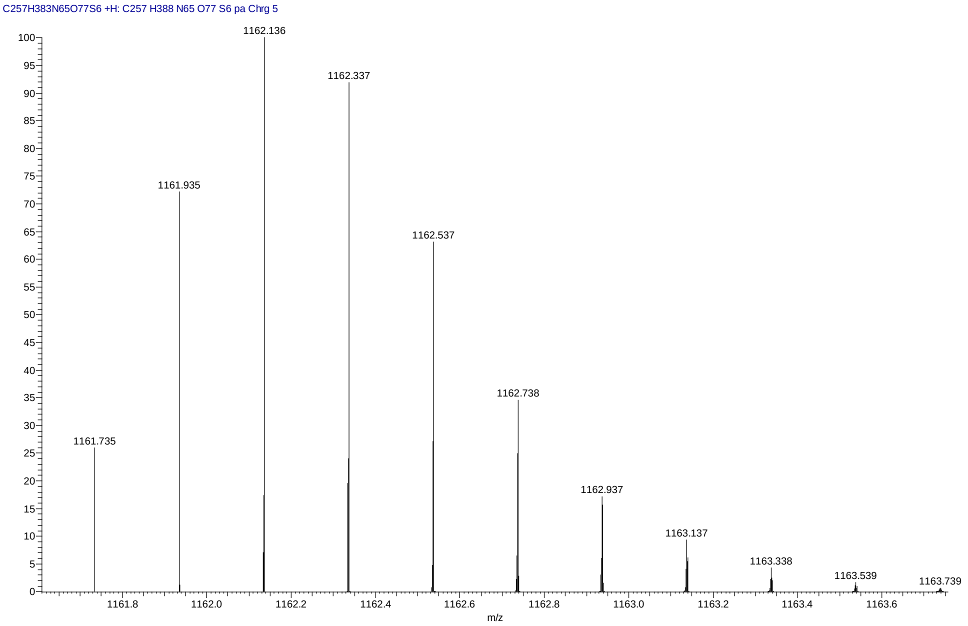

### Insulin A-chain

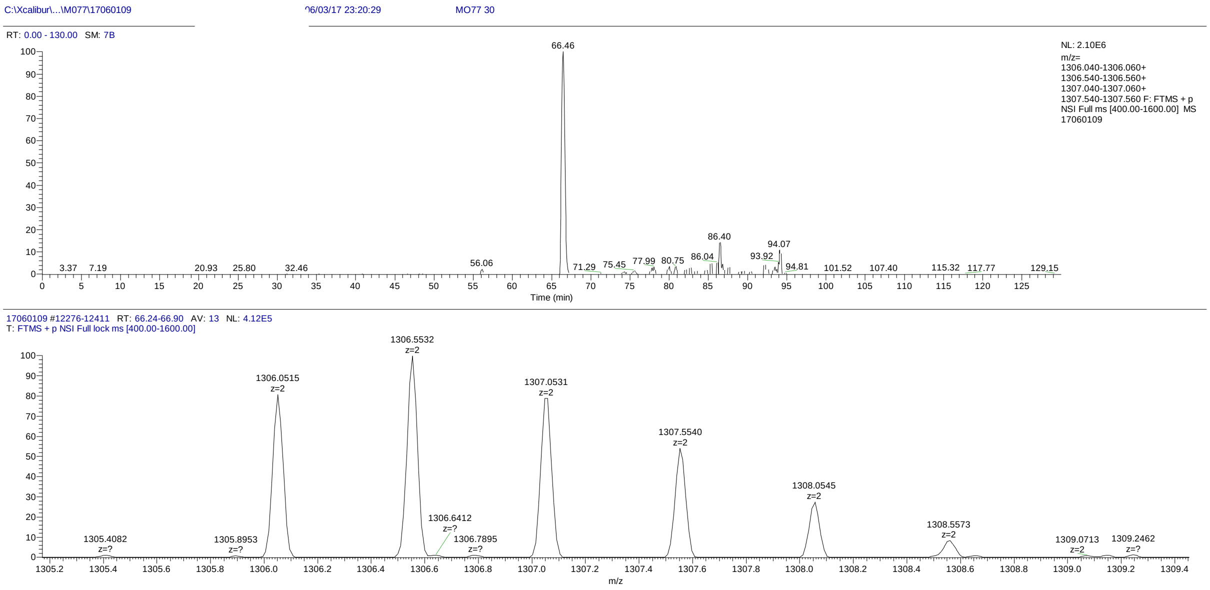


### Theoretical isotope pattern for $[M+2H]^{2+}$ charge state

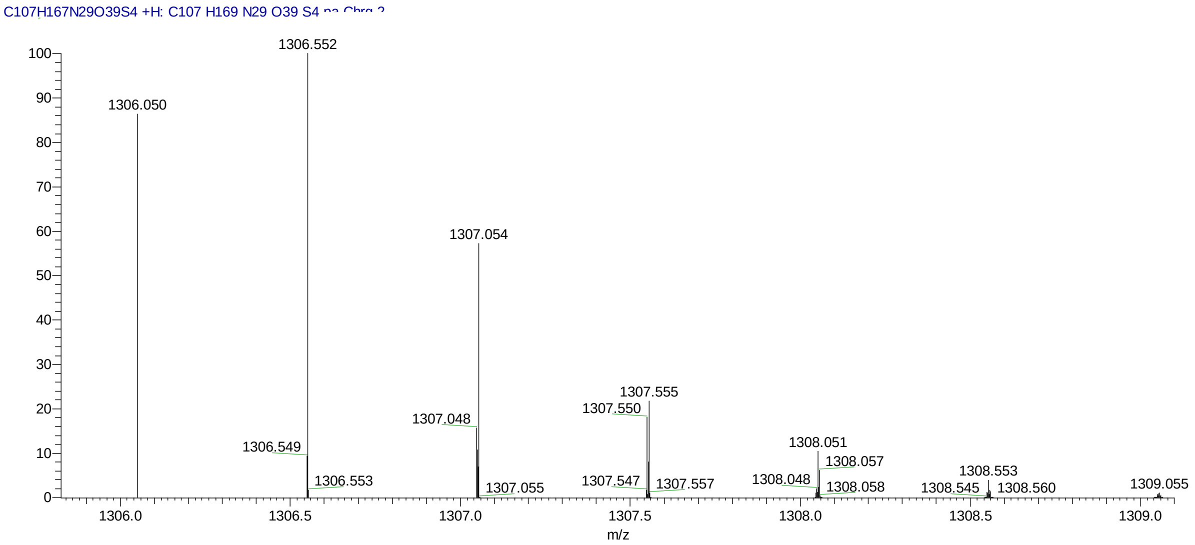

### Insulin B-chain

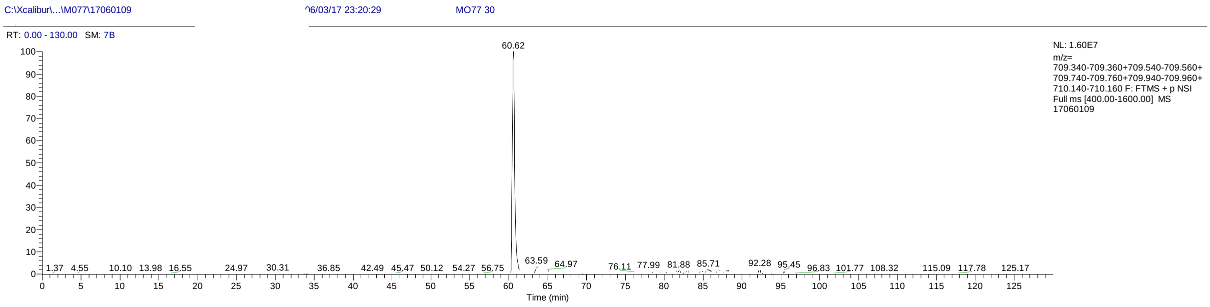


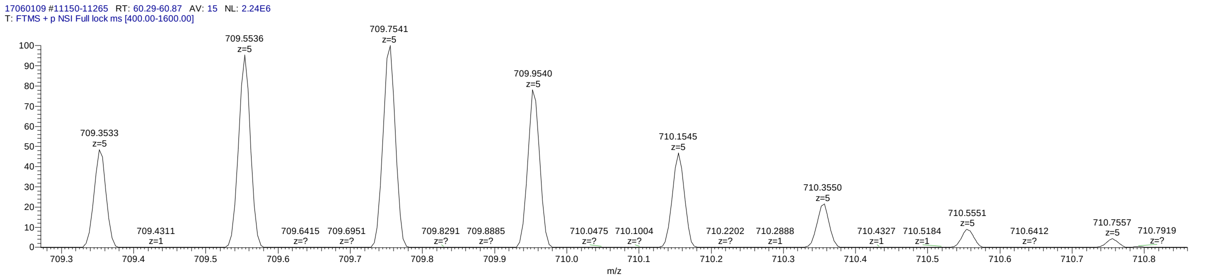


### Theoretical isotope pattern for $[M+5H]^{5+}$ charge state

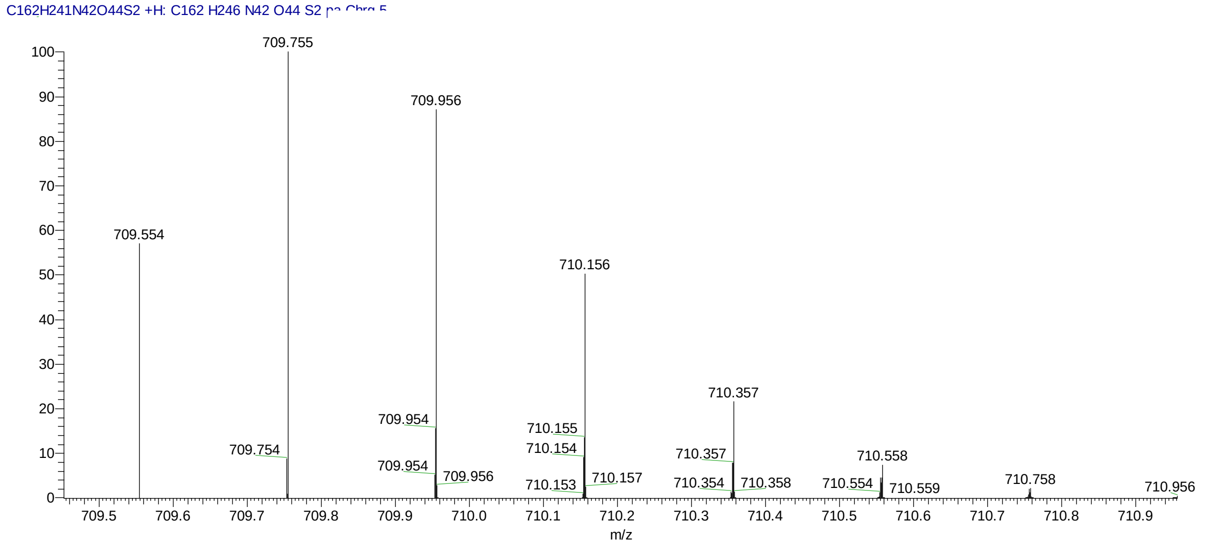

### Insulin C-peptide

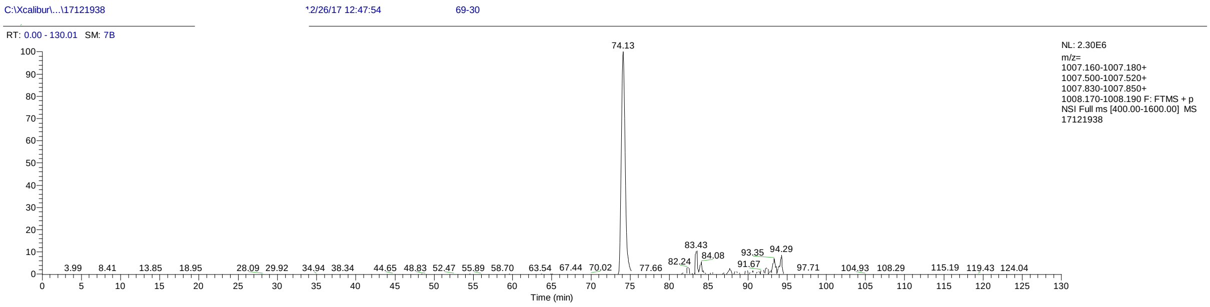


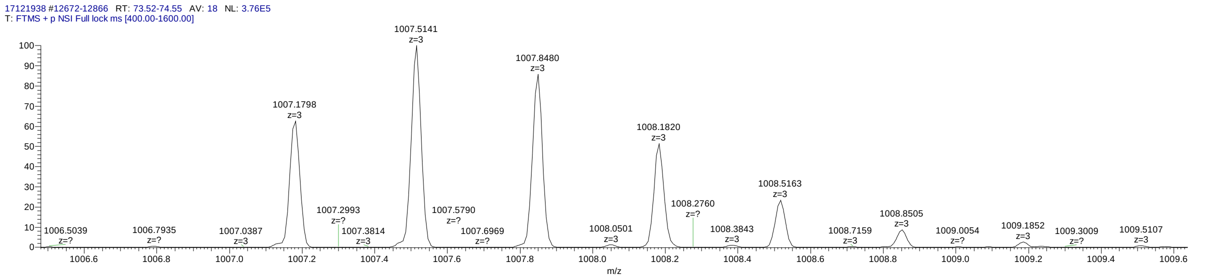


### Theoretical isotope pattern for $[M+3H]^{3+}$ charge state

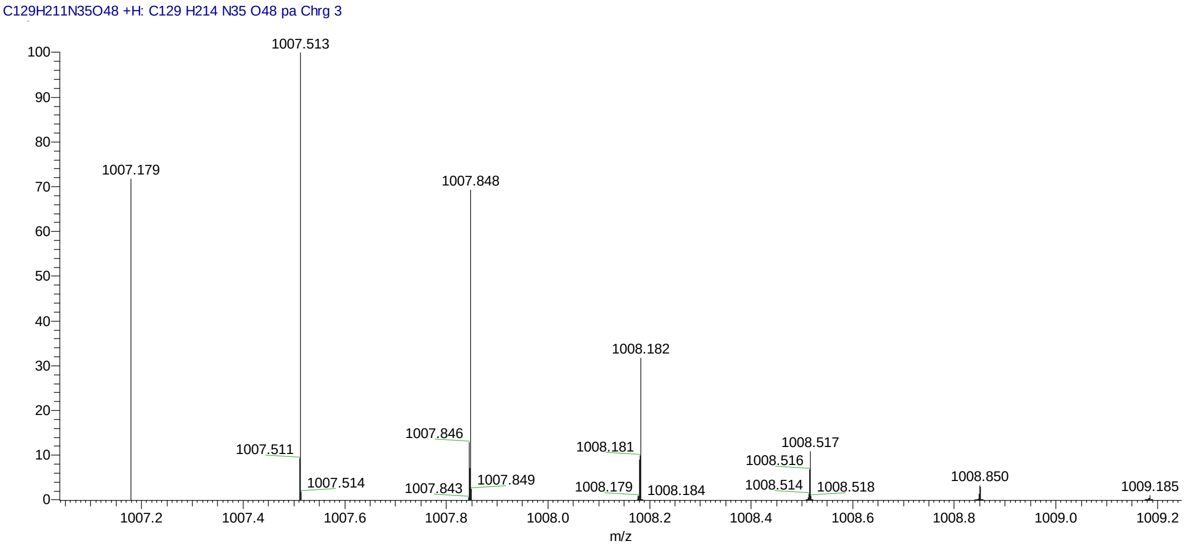

### Bovine insulin (intact)

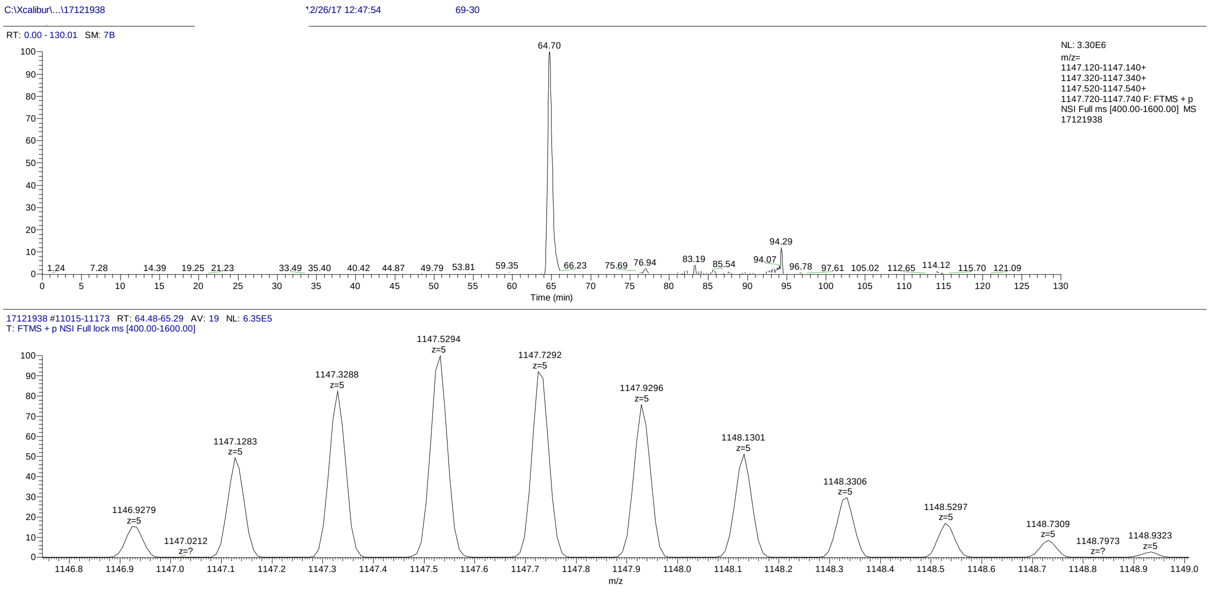


### Theoretical isotope pattern for $[M+5H]^{5+}$ charge state

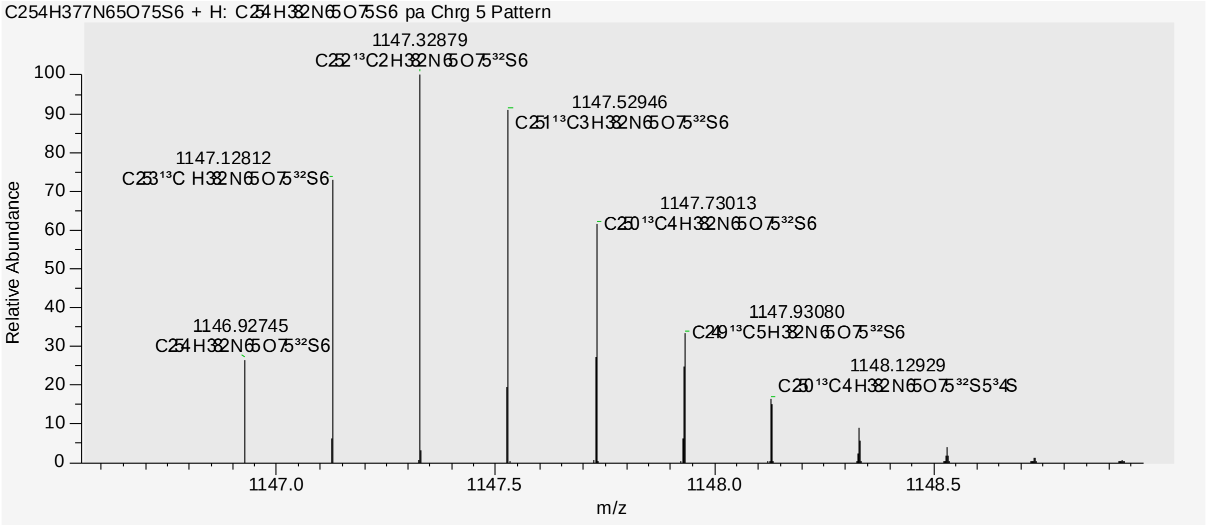

### GIP propeptide

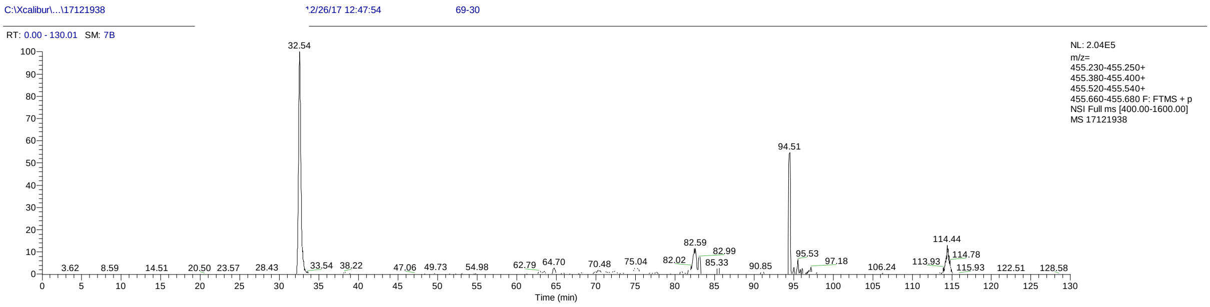


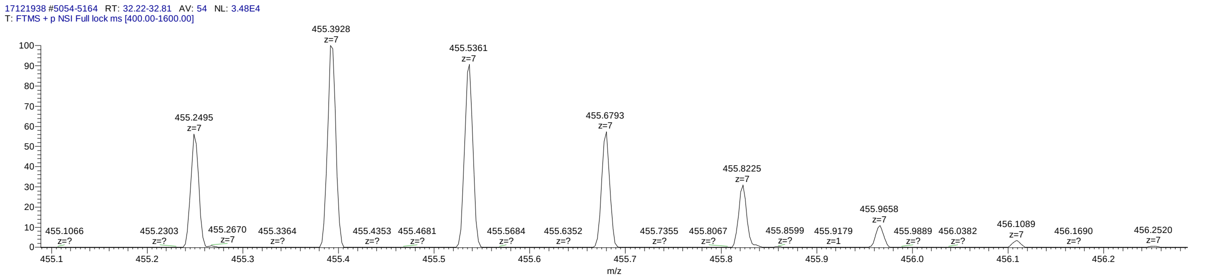


### Theoretical isotope pattern for $[M+7H]^{7+}$ charge state

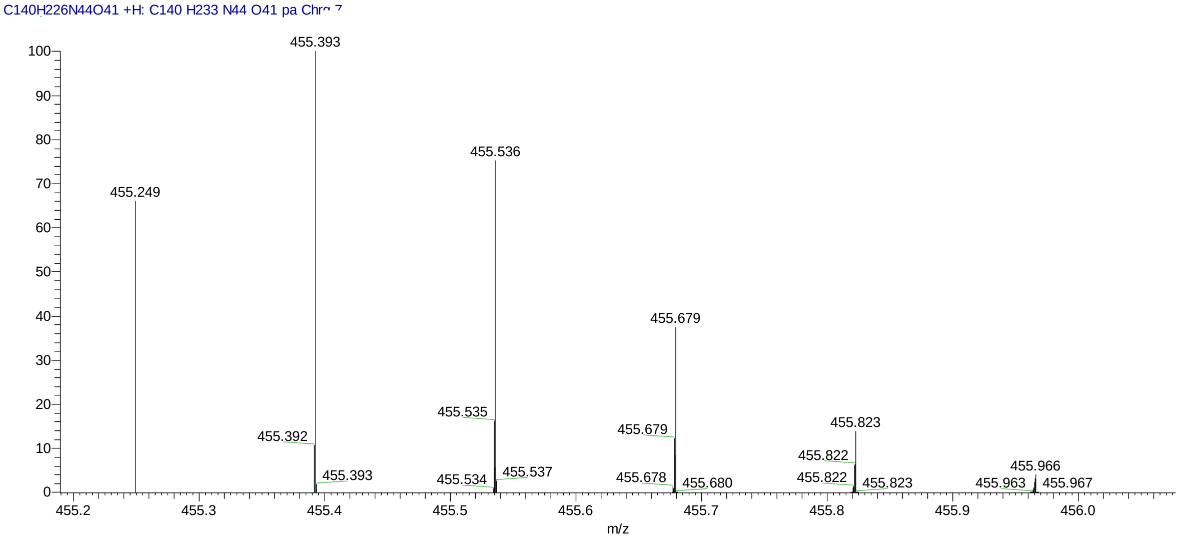

### GIP 1-42

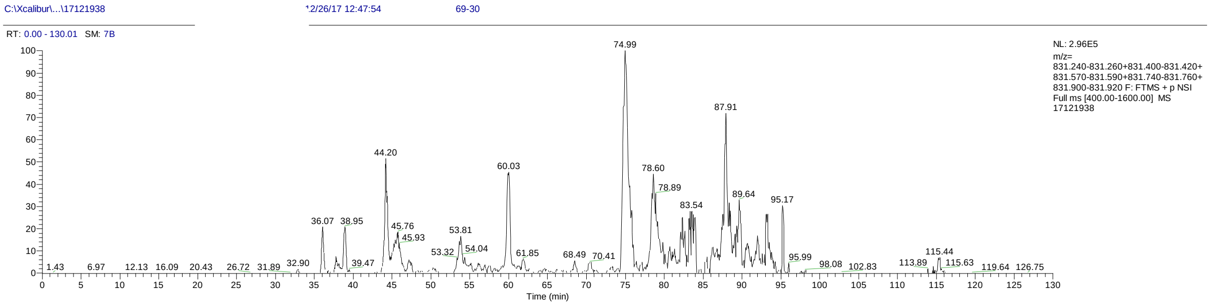


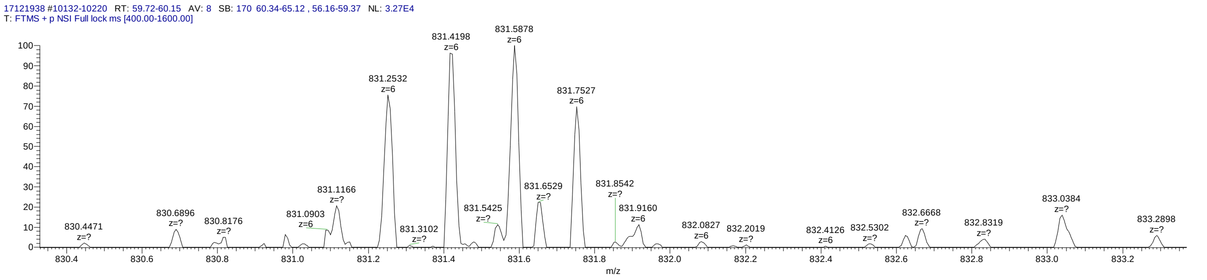


### Theoretical isotope pattern for $[M+6H]^{6+}$ charge state

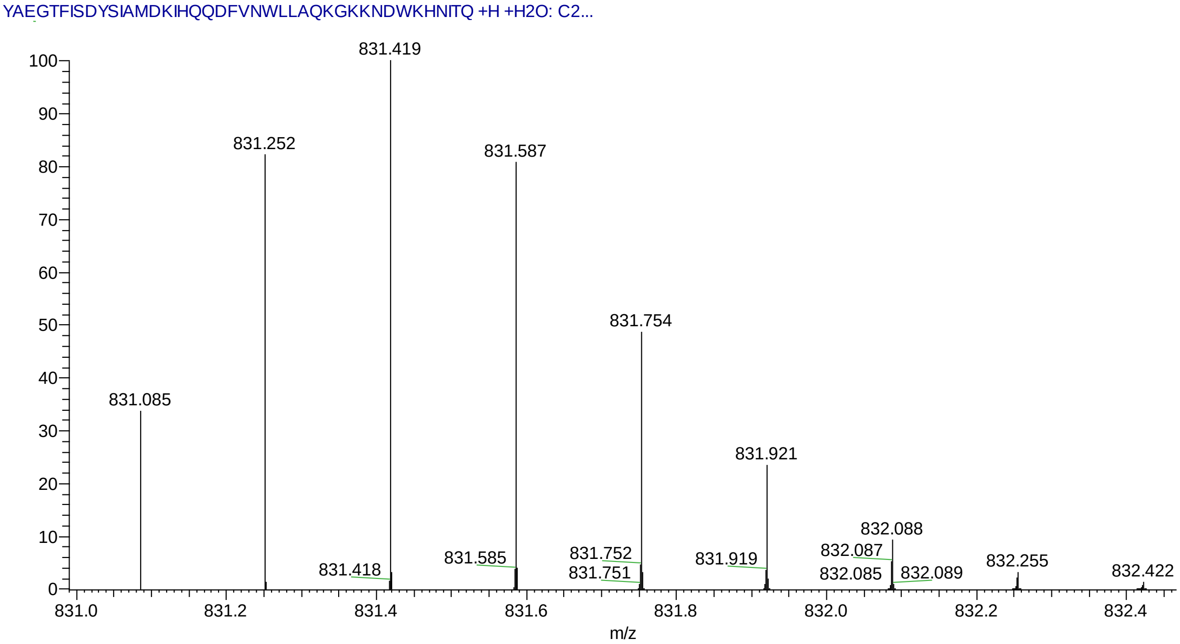

### GIP 3-42

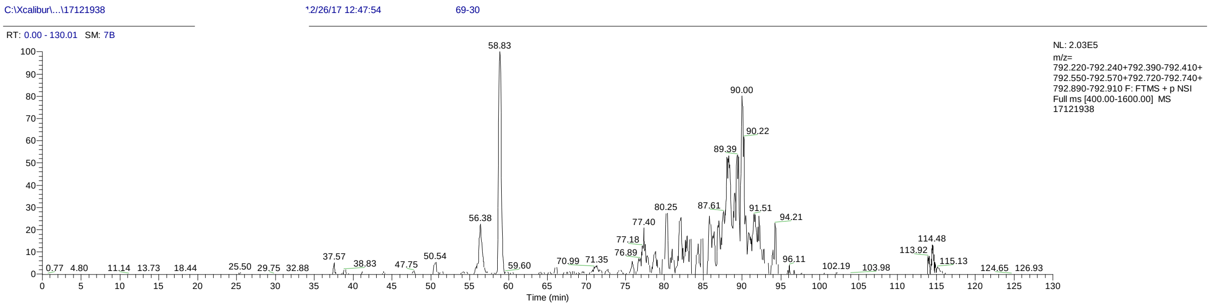


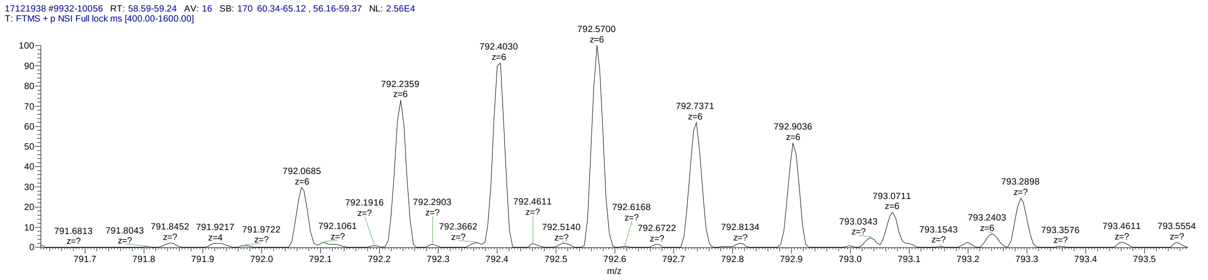


#### Theoretical isotope pattern for $[M+6H]^{6+}$ charge state

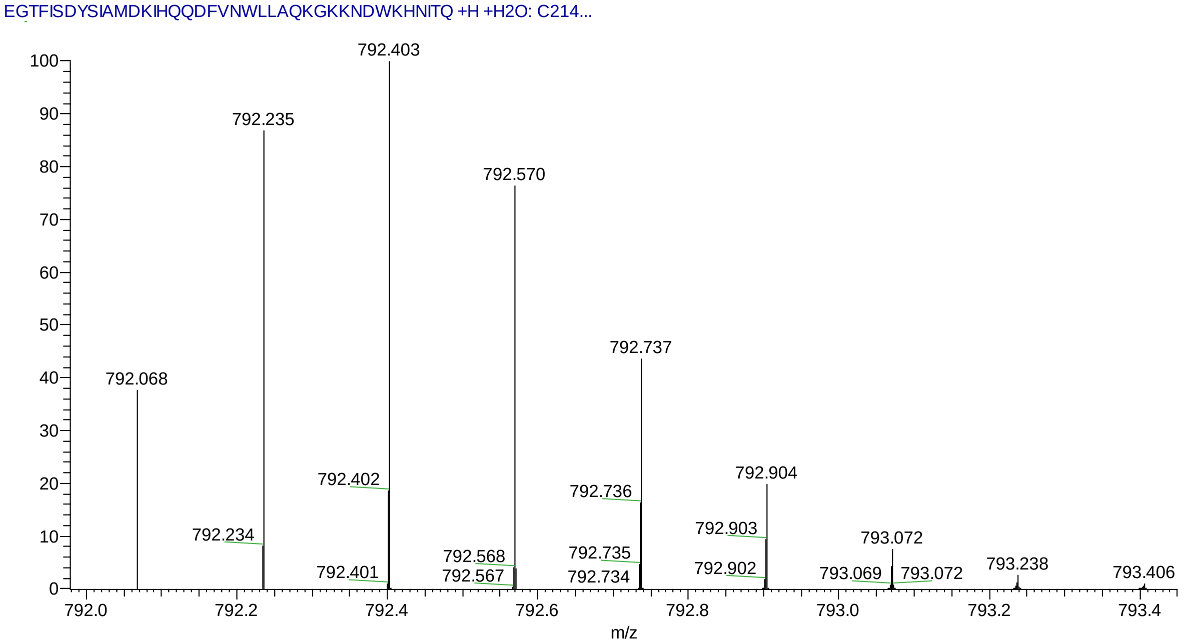

### Neurotensin

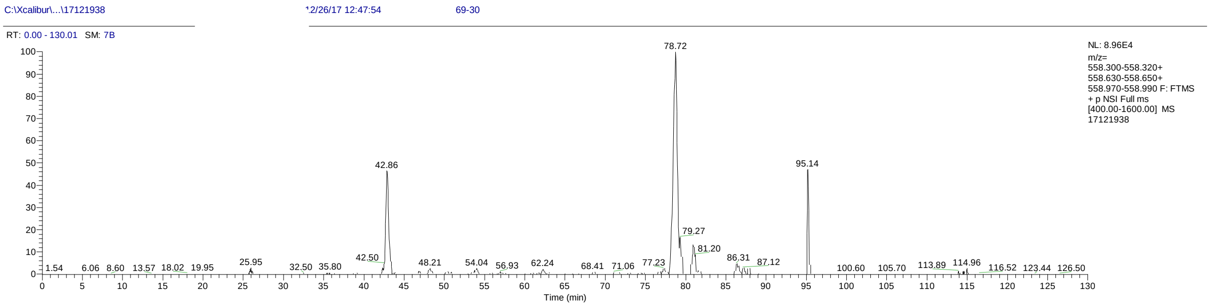


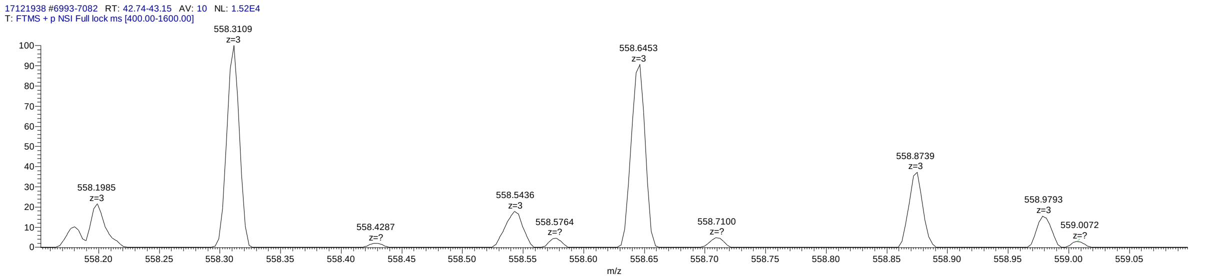


### Theoretical isotope pattern for $[M+3H]^{3+}$ charge state

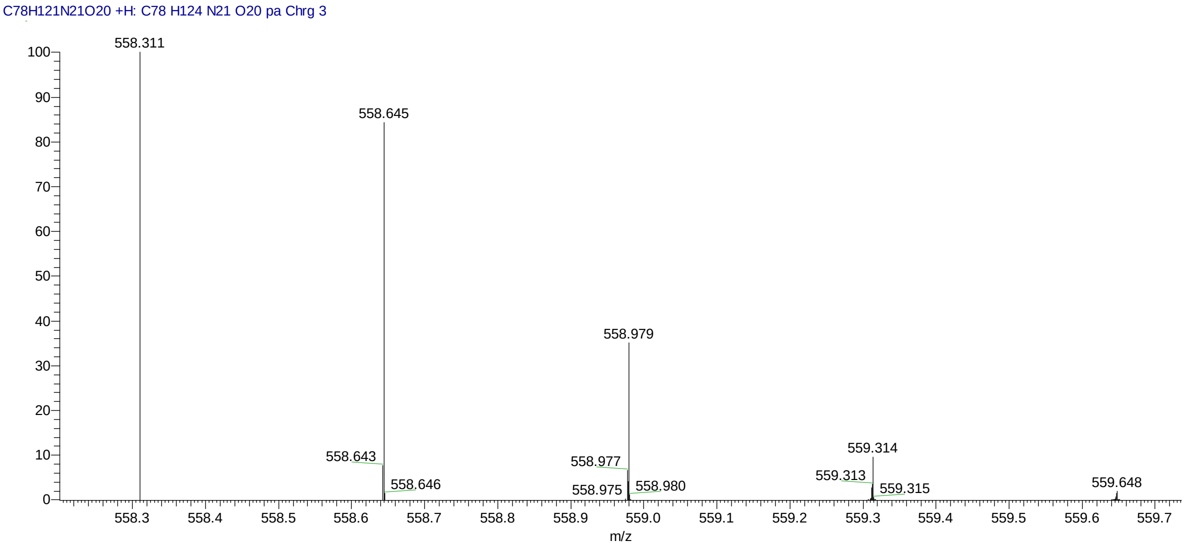

### Motilin

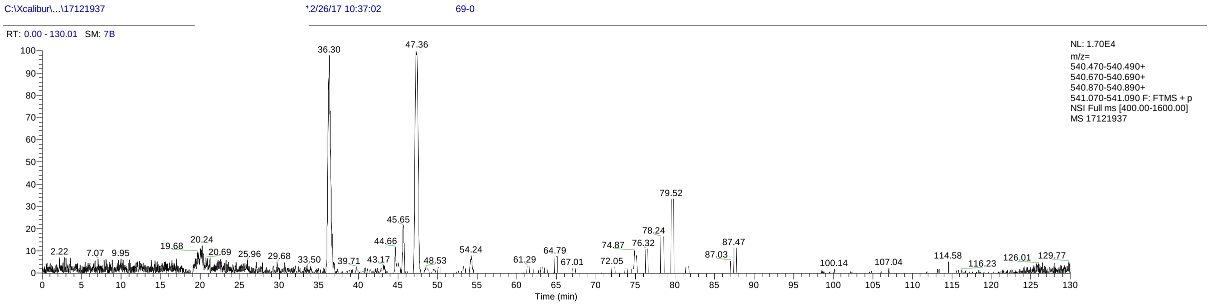


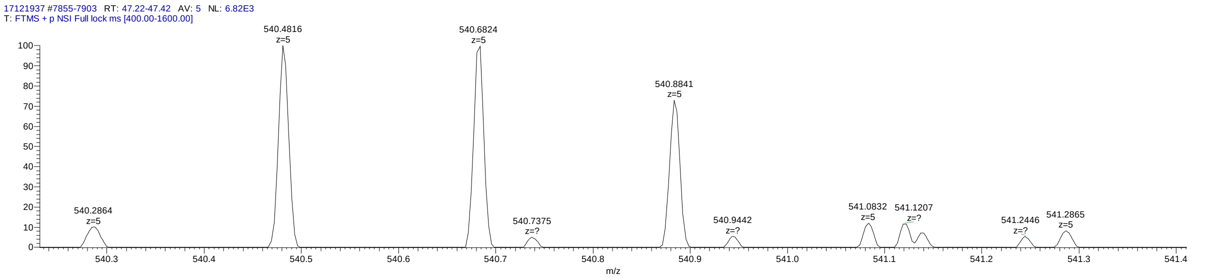


### Theoretical isotope pattern for $[M+5H]^{5+}$ charge state

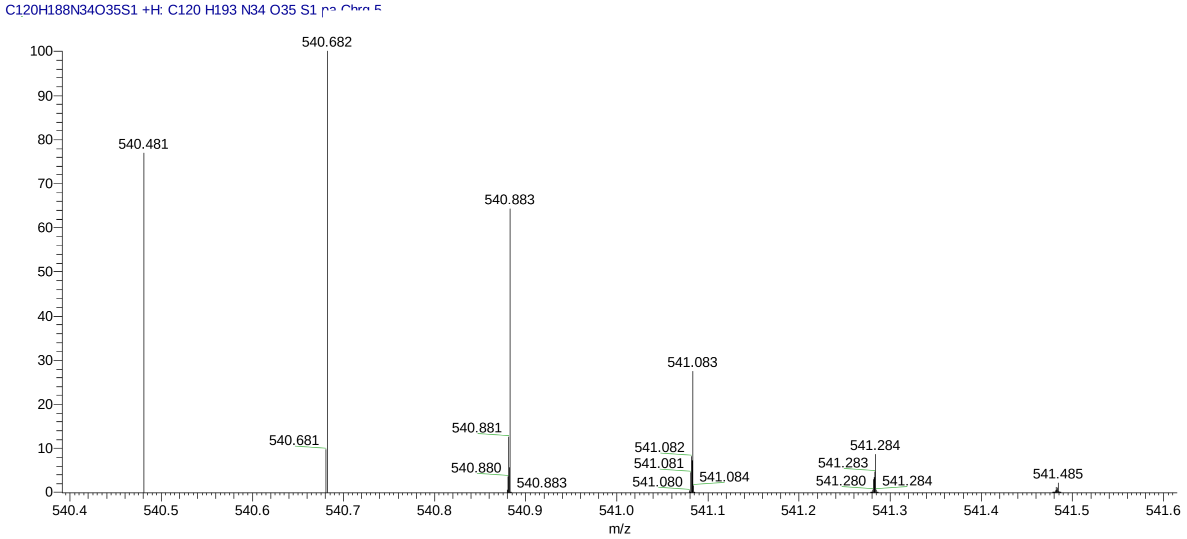

### Adrenomedullin (45-92)

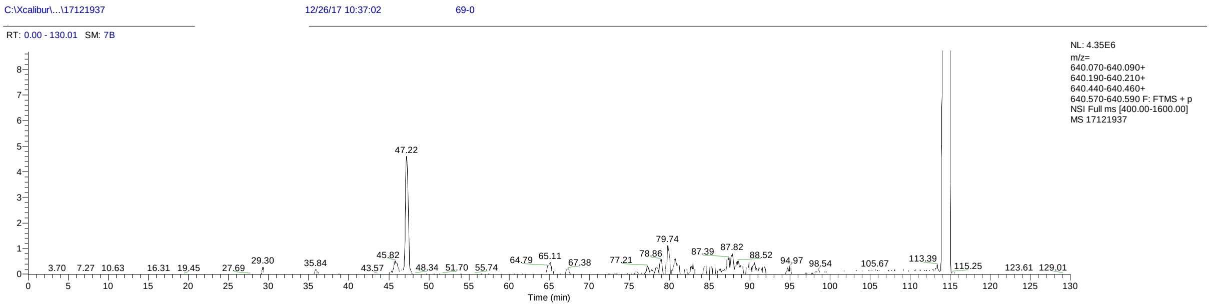


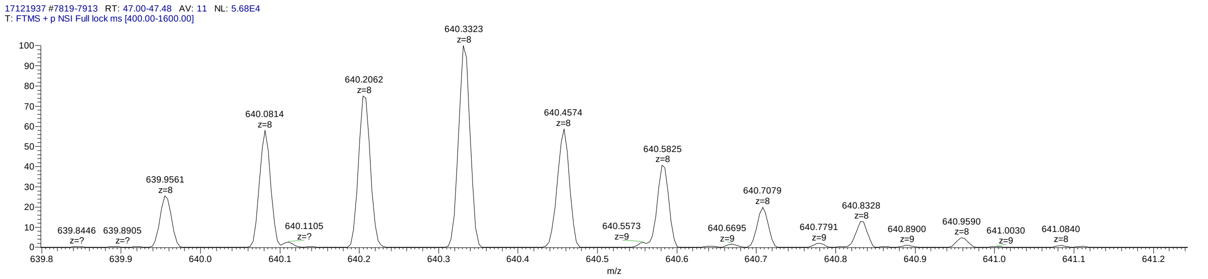


### Theoretical isotope pattern for $[M+8H]^{8+}$ charge state

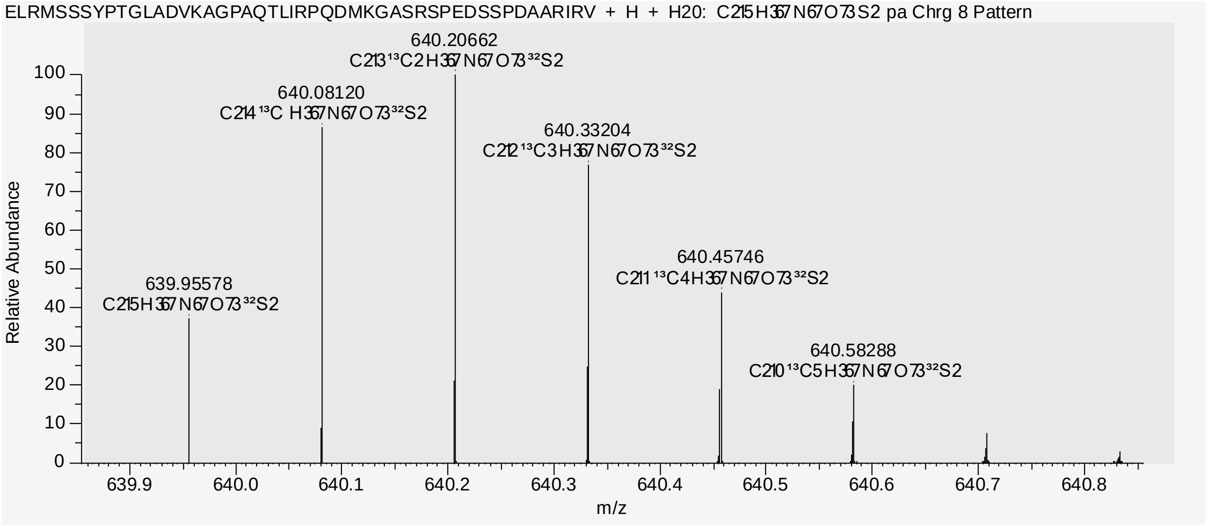

#### Augurin (45-92)

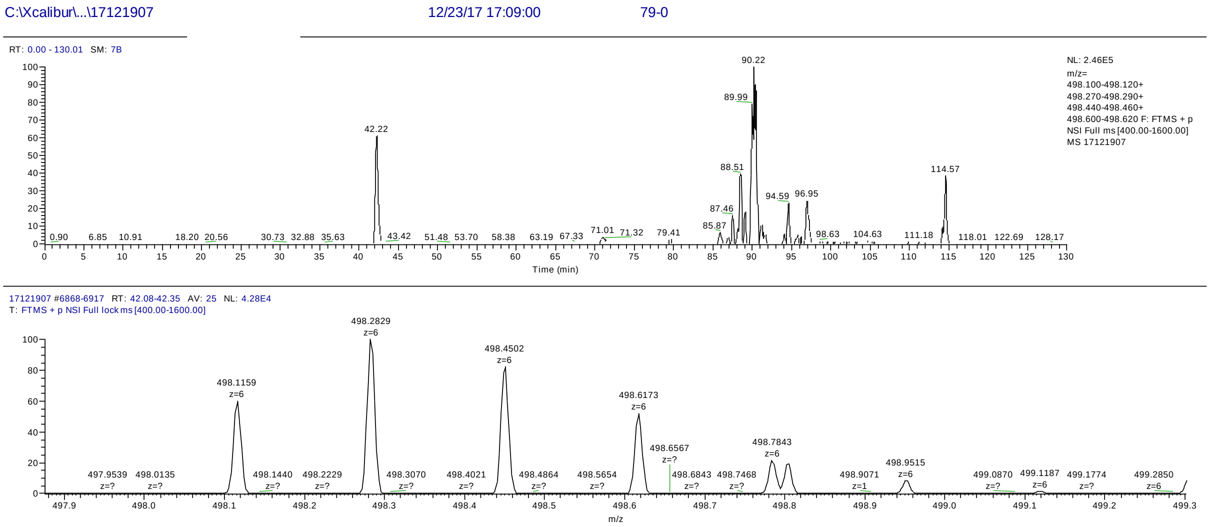


#### Theoretical isotope pattern for $[M+5H]^{5+}$ charge state

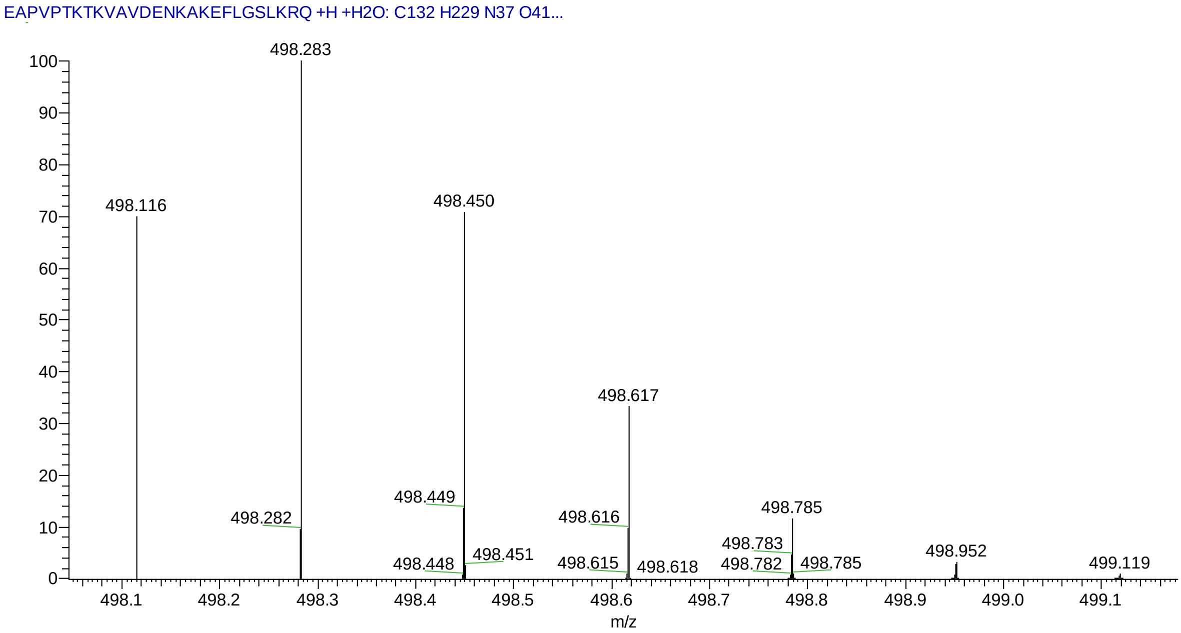
